# Allelic correlation is a marker of tradeoffs between barriers to transmission of expression variability and signal responsiveness in genetic networks

**DOI:** 10.1101/2021.11.26.470134

**Authors:** Ryan H. Boe, Vinay Ayyappan, Lea Schuh, Arjun Raj

**Affiliations:** Genetics and Epigenetics, Cell and Molecular Biology Graduate Group, Perelman School of Medicine, University of Pennsylvania, Philadelphia, PA, USA; Department of Bioengineering, School of Engineering and Applied Sciences, University of Pennsylvania, Philadelphia, PA, USA; Institute of AI for Health, Helmholtz Zentrum München - German Research Center for Environmental Health, 85764 Neuherberg, Germany; Department of Mathematics, Technical University of Munich, Garching, 85748, Germany; Department of Genetics, Perelman School of Medicine, University of Pennsylvania, Philadelphia, PA, USA

## Abstract

Accurately functioning genetic networks should be responsive to signals but prevent transmission of stochastic bursts of expression. Existing data in mammalian cells suggests that such transcriptional “noise” is transmitted by some genes and not others, suggesting that noise transmission is tunable, perhaps at the expense of other signal processing capabilities.However, systematic claims about noise transmission in genetic networks have been limited by the inability to directly measure noise transmission. Here we build a mathematical framework capable of modeling allelic correlation and noise transmission. We find that allelic correlation and noise transmission correspond across a broad range of model parameters and network architectures. We further find that limiting noise transmission comes with the trade-off of being unresponsive to signals, and that within the parameter regimes that are responsive to signals, there is a further trade-off between response time and basal noise transmission. Using a published allele specific single cell RNA-sequencing dataset, we found that genes with high allelic odds ratios are enriched for cell-type specific functions, and that within multiple signaling pathways, factors which are upstream in the pathway have higher allelic odds ratios than downstream factors. Overall, our findings suggest that some degree of noise transmission is required to be responsive to signals, but that minimization of noise transmission can be accomplished by trading-off for a slower response time.

## INTRODUCTION

Cells must balance stability of their internal workings and responsiveness to external changes. One way in which such information processing is accomplished is via genetic networks, in which the products of certain genes regulate the expression of other genes (or themselves). A core function of these genetic networks is to respond to and process signals and translate them into changes in gene expression. However, genetic networks are also subject to random fluctuations in the numbers and activity of the constituent molecules (i.e., molecular “noise”) (Becskei and Serrano, 2000; Elowitz et al., 2002; Løbner-Olesen, 1999; Ozbudak et al., 2002; Raj and van Oudenaarden, 2008; Raser and O’Shea, 2005; Symmons and Raj, 2016). Such noise threatens the accurate functioning of genetic networks because, if transmitted and amplified through the network, noise may mimic the presence of a signal even when no such signal exists. In human cells, the few existing studies show that noise transmits in some contexts but not others, suggesting that the degree of noise transmission of a genetic network may be an important, tunable feature (Jena et al., 2021; Shah and Tyagi, 2013). What remains unclear is what the consequences of suppressing noise transmission may be; in particular, what other signal processing limitations must networks have to achieve such noise suppression?

The current data in human cells suggests that some systems limit noise transmission while others do not, but the scarcity of data makes systematic conclusions difficult to make. Consequently, it is hard to say which genetic networks either allow or block noise transmission and what sorts of tradeoffs may arise from those constraints. For instance, one potential tradeoff could be that increased noise transmission is the cost of having an ability to respond to dynamic signals. An illustration of such a potential tradeoff is the expression of the immediately early genes *FOS* and *JUN*. In serum-starved HeLa cells, serum stimulation induced noisy and uncorrelated transcription of *C-FOS* and *C-JUN* that was buffered by heterodimerization of their protein products, preventing more noise in two of the downstream targets (Shah and Tyagi, 2013). However, in mouse epidermis, *Fos, Jun*, and other immediate response genes showed correlated expression in single cells, suggesting that upstream variability was consistently transmitted downstream to multiple targets in the same cell (Jena et al., 2021). Notably, in mouse epidermis, the *Mapk1, Fos, Jun* pathway is an important developmental signal, controlling basal stem cell and keratinocyte differentiation (Hiratsuka et al., 2020; Mehic et al., 2005). Thus, in an active, developing context, the Fos-Jun pathway may exhibit high noise transmission, but in the resting context of *in vitro* HeLa cells, the same pathway blocks transmission of noise. This example raises the possibility that the design and parameters of genetic networks may be tweaked to make different tradeoffs when faced with different functional requirements.

It is currently difficult to directly measure noise transmission due to lack of experimental tools, so in order to circumvent this limitation, we sought to develop a more easily-measurable proxy for noise-transmission. Presently, most available data consists of either static measurements of endogenous RNA; real time measurements of multiple RNA or protein species in the same pathway would allow direct measurements of noise transmission, but this technology is just emerging and is technically challenging (Cohen et al., 2008; Frenkel-Morgenstern et al., 2010; Wan et al., 2021; Zimmer et al., 2017).The limited experimental data measuring noise transmission has led to a lack of models describing noise transmission and its consequences. We thus sought to build a framework that could model noise transmission while also being easily related to ample and readily accessible experimental data. Such a framework would not only circumvent experimental challenges but also allow us to re-examine existing data for evidence of signal processing tradeoffs that may arise from limiting noise transmission.

Can the correlation between expression of two copies of a gene over time serve as an indicator for noise transmission (Figure 1A)? Though previously considered for two copies of a gene in *E*.*coli* (Elowitz et al., 2002), we propose that the degree of expression correlation between two endogenous alleles of a gene similarly can be a proxy for noise transmission. For an intuition as to how allelic correlation is related to noise transmission, consider an upstream regulator that controls expression of two alleles of a gene equally. With a constant low level of regulator, each allele will burst independently, leading to low allelic correlation. With a constant high level of regulator, each allele will independently burst albeit at a higher rate. However, with a transient spike in the upstream regulator, will both alleles simultaneously burst? If so, then the variability has been transmitted from the upstream regulator to both downstream alleles, leading to high allelic correlation. The transmission of variability that results in correlated alleles is conceptually similar to noise transmission and is also measured by single cell RNA sequencing or RNA FISH (Deng et al., 2014; Levesque and Raj, 2013; Levesque et al., 2013; Reinius et al., 2016; Symmons et al., 2019). Should the correspondence between allelic correlation and noise transmission hold, mathematical models that include both measurements could be tuned to fit available data and also could generate testable hypotheses.

**Figure 1.**
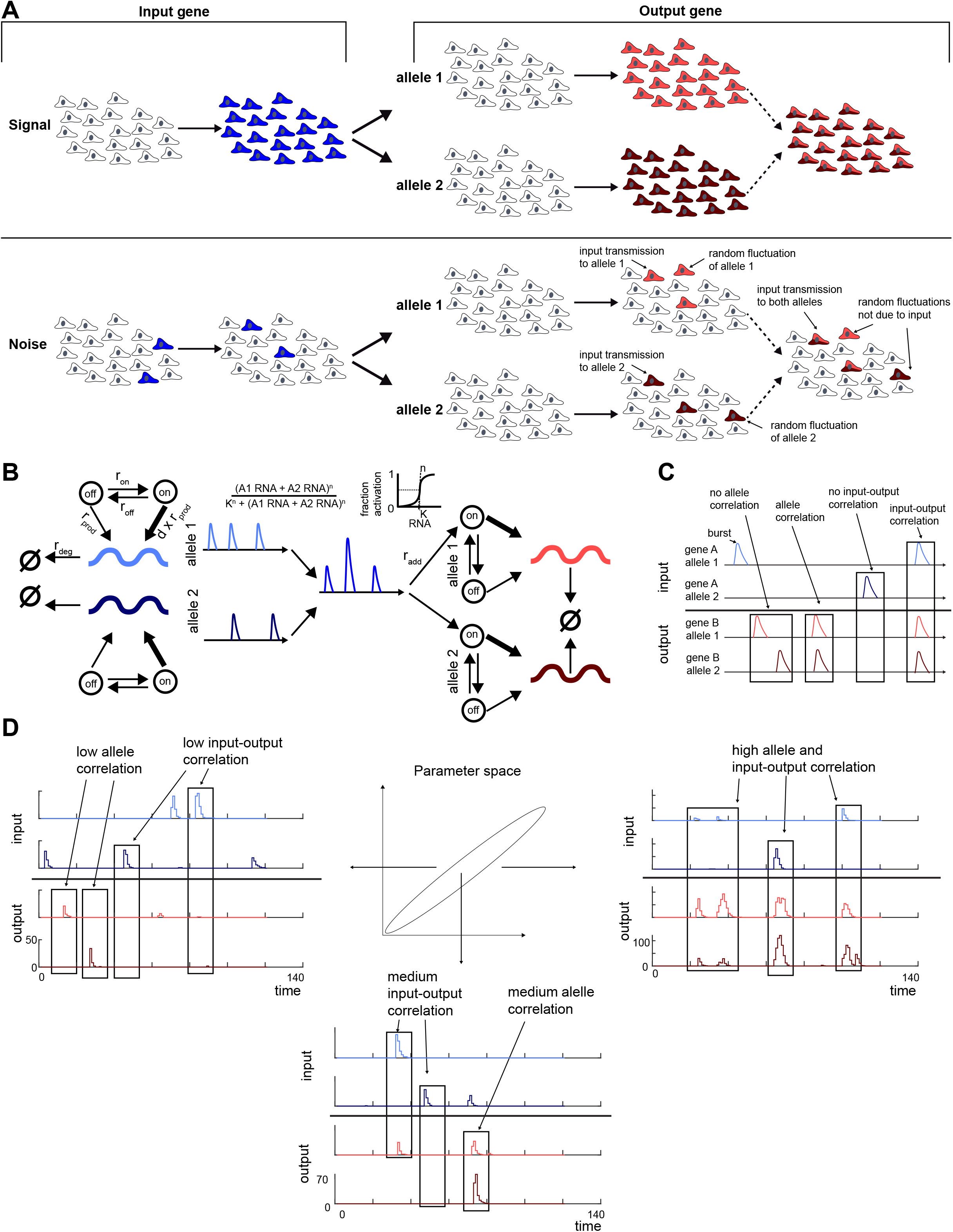
An allele-specific transcriptional bursting-based network model can be tuned to either block or transmit noise, as captured by the degree of allelic expression correlation. (A) Schematic of a population of cells responding to either an upstream signal (top) or noise (bottom). When some cue causes an upstream gene to burst (top, left), that signal needs to be transmitted to the downstream gene to cause it to burst (top, right). When there is no cue and there are random bursts in the upstream node (bottom, left), cells in which the bursts in the upstream genes occur may have activation of the downstream genes (bottom, right). This noise may transmit to both alleles of the downstream gene, causing co-expression of the alleles. (B) Schematic of allele-specific transcriptional bursting-based network model for example two node network. Gene A and gene B both are modeled as having two identical alleles with DNA in either an inactive (off) or activate (on) state, with transitions governed by rates r_on_ and r_off_. Each allele of the gene produces distinct mRNA, synthesized with rates r_prod_ and d*r_prod_ in the inactive and active states respectively. mRNA from each allele degrades with rate r_deg_. Gene regulation is modeled by a Hill function using the sum of mRNA counts from both alleles of a regulating gene as an input, governed by Hill coefficient n, dissociation constant k, and rate r_add_. Regulation is applied equally to the DNA of both alleles. (C) Schematic of examples of allelic correlation and input-output correlation for two genes with two alleles each. Allele correlation is when both alleles burst at the same time. Input-output correlation is when the bursting of either allele of a gene causes bursting of its regulated gene. (D) Schematic of parameter space with real example traces of a two node, two allele network with gene A regulating gene B. Depending on the parameters chosen, simulations of the network show low or high allelic and input-output correlation. We observe that allelic correlation qualitatively corresponds to input-output correlation.

Here we describe a mathematical framework to test the correspondence between allelic correlation and noise transmission and to model the signal processing tradeoffs that may result from limiting noise transmission. We simulated networks of varying size and connectivity defined by a large range of parameters to find that in this minimal model, allelic correlation and noise transmission are tightly coupled across a broad range of parameters and network architectures. We further found that response to external signals was only possible in simulations that also exhibited high noise transmission, but that among simulations that could respond to signals, there simulations could trade a slower signal response time for less basal noise transmission. To test these predictions, we looked at an allele specific single cell RNA-sequencing dataset and found that upstream members of three signaling pathways had high allelic odds ratios in agreement with the results from our model. Overall, we demonstrated that noise transmission could be inferred by allelic correlation and that limiting noise transmission in genetic networks led to the trade-off of being unresponsive to external signals.

## RESULTS

### Selection of mathematical framework to model both correlated allelic expression and noise transmission

#### Description of mathematical model

Our goal was to measure noise transmission in mathematical models of genetic networks and systematically describe the tradeoffs in signal response that arise from reducing noise transmission. However, noise transmission remains difficult to measure directly from experiments. We sought out a proxy for noise transmission that is readily measured experimentally and thus would allow us to use more experimental data. Recent advances in RNA FISH and single cell RNA sequencing have made correlation between alleles in single cells easier to measure, and so we wanted to establish whether there was a systematic quantitative relationship between allelic correlation and noise transmission which we could exploit. In order to do so, we needed to simultaneously model allelic correlation and noise transmission across broad swaths of parameters and in many configurations of genetic networks.

We wanted to construct a model with the minimal components needed to recapitulate allelic correlation, transcriptional noise transmission, and transcriptional response to signals. Because promoter leak can modulate expression noise in certain cases (Huang et al., 2015; Venturelli et al., 2012), we used a leaky telegraph model as a starting point for our model (Figure 1B). In the inactive state, the promoter produces gene product at a very low “leak” rate but can stochastically enter the active state to produce a large “burst” of gene product. These transcriptional bursts are one critical means by which variability is created (Raj et al., 2006). We took this well-known model of transcription (Ham et al., 2020; Kepler and Elston, 2001; Peccoud and Ycart, 1995; Raj et al., 2006; Schuh et al., 2020; Thomas et al., 2014) and used it as the basis for modeling gene network interactions. Each gene in the network has two alleles, each with identical but independent chemical reactions and rates. Each allele produces a gene product that is computationally distinguishable but is equivalent in terms of functionality in the model; i.e., both are equally capable of binding a downstream promoter and affecting consequent gene expression. The chemical reactions of our model include reversible switching of the gene between an active (r_on_) and inactive (r_off_) state, with an additional rate of switching to the active state from promoter binding from an upstream regulator (r_add_). r_add_ depends on the total amount of gene product of both alleles of the regulating node to which we applied a Hill function, which includes additional parameters: Hill coefficient n (to capture the cooperativity of the interaction) and dissociation constant k (to capture the affinity of the transcription factor). When the gene is in the inactive state, the gene is transcribed as a Poisson process at a very low basal rate (r_prod_), and when the gene is active, the rate becomes higher by a constant factor (d, where d ≥ 1) for a total of d*r_prod_. We consider degradation of RNA as a Poisson process with rate r_deg_. Overall, our model has a total of eight independent parameters per allele (see methods for table summarizing the parameters). For simplicity, we assumed all alleles to be governed by the same parameters.

#### Definition of metrics and initial characterization of model

We expected that the above model would be able to capture ranges of allelic correlation and noise transmission. We could use the time traces of simulations of our model to see whether our model could produce different values of allelic correlation and noise transmission. Spontaneous bursting of gene expression in our model is considered “noise,” and thus noise transmission is when a burst in expression of the upstream gene triggers the bursting of a downstream gene (Figure 1C). We refer to this co-expression of upstream and downstream genes as input-output correlation. In contrast, we defined allelic correlation to be when both alleles of the same gene burst at the same time. Importantly, these two phenomena need not be inherently coupled, since the promoter and gene product of each allele is a separate species with a separate chemical reaction in our model. Our model extended a multi-node leaky-telegraph network model of transcription to include multiple alleles, thus permitting simultaneous measurement of allelic correlation and noise transmission.

To establish that we were able to differentiate high and low allelic and input-output correlations and to qualitatively assess which parameters lead to each situation, we used Gillespie’s algorithm (Gillespie, 1977) to simulate the simplest possible network of a single upstream gene connected to a single downstream gene. We then visually inspected time traces of simulations across parameters (Figure 1D). By manual inspection of the time traces, we found instances of parameters for which both allelic and input-output correlation were low, i.e., alleles rarely burst together and upstream bursts were not transmitted downstream (Figure 1D, left plots). We also found parameter sets for which allelic and input-output correlation were high, i.e., bursts were always transmitted downstream and both alleles always burst together (Figure 1D, right plots). Finally, we found a small set of parameters with seemingly intermediate allelic and input-output correlation, in which an upstream burst would often transmit downstream, but only triggered a single allele (Figure 1D, bottom plots). These individual observations confirmed that our model could be tuned to both transmit and block transmission of noise. We conclude that, at least in the small number of parameter sets we could visually inspect, allelic correlation can serve as a proxy for input-output correlation.

To assess the generality of any findings, we aimed to test networks of varying size and structure with a large number of parameter sets. We wanted to find network configurations that admitted high and low allelic and input-output correlation and test the effects of network size and connectivity. In our model, nodes represented genes and directed edges represented regulating interactions between genes. The connectivity, or number of ingoing edges for any node in the network, indicates the number of regulating interactions for each gene. For the sake of simplicity and computational tractability, we used symmetric networks, which capture most network interactions without loss of generality but have a far more limited set of parameters (Schuh et al., 2020). We also only analyzed networks of distinct isomorphism classes because our assumption of symmetric behavior means that isomorphic networks are formally equivalent. We additionally excluded networks that were compositions of smaller subnetworks to prevent re-analyzing networks. We chose to highlight results from a five node connectivity one network, although we also ran simulations with other numbers of nodes and degrees of connectivities (Figure S1).

### Allelic odds ratio captures high correlated allelic expression as a ridge in r_add_/r_off_ parameter space

Though we saw low and high allelic correlation in a limited number of parameter sets, we wanted to systematically determine which parameter values led to high and low allelic correlation. To do so, we first needed a single metric to capture the degree of allelic correlation in a given simulation. To that end, we developed an algorithm to calculate the odds ratio of simultaneous expression of both alleles of a given gene (Figure 2A). We first selected a threshold value and binarized the entire time series simulation data. We used a threshold of three molecules to exclude instances of “promoter leak” from an inactive promoter that result in production of a small number of RNA molecules. After binarizing both alleles of a gene, we summarized whether neither, one, or both alleles were on at each timepoint in a contingency table. Finally, we computed the odds ratio for that gene. Since all networks were symmetric, the odds ratios should in principle be equal for each gene in a given simulation: therefore, for ease of visualization and analysis, we used the mean odds ratio across all genes in a given simulation. An odds ratio of greater than one indicates that alleles express at the same time more often than expected if they were independent processes, an odds ratio of one means the alleles are expressed at the same time no more than expected by chance, and an odds ratio of less than one indicates that the alleles are expressing less than expected by chance, which is usually interpreted as anticorrelation.

**Figure 2.**
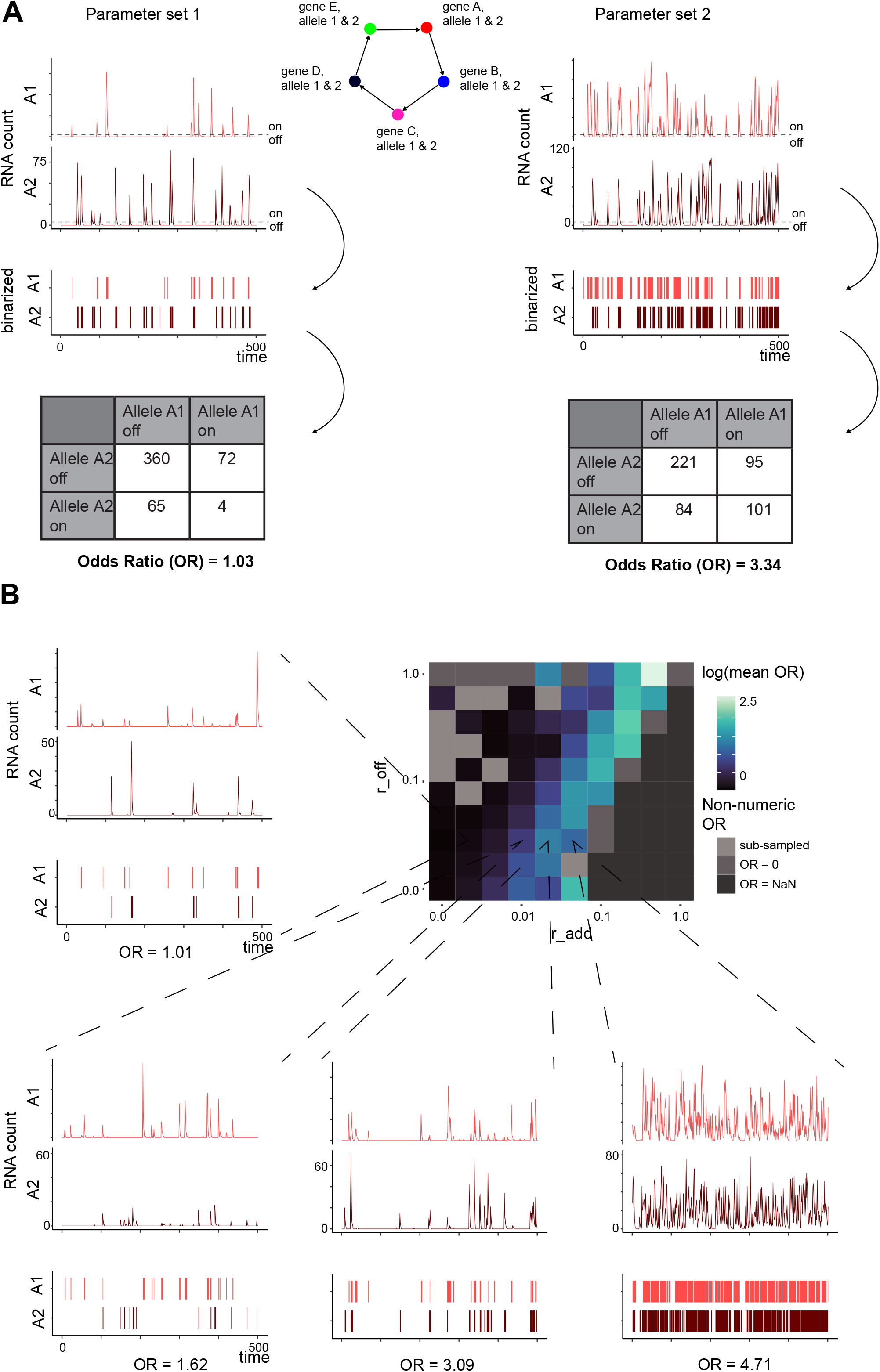
Odds ratio for allelic expression shows a ridge of high allelic correlation in the parameter space defined by r_add_ and r_off_. (B) For each gene in a simulation, we set a threshold of 3 molecules and binarize the RNA count over time for each allele. At each time point, we are interested if neither, only one, or both alleles are expressed. We calculate the contingency table using each time point as an observation. We then calculate the odds ratio where a higher odds ratio means more co-expression of the alleles and an odds ratio of one is no more co-expression than by chance. Finally, we calculate the mean odds ratio across all genes in a given simulation. (B) A heatmap showing the distribution of log transformed mean odds ratio in the parameter space defined by rates r_add_ and r_off_. Other parameters are held constant at k=110, r_on_ = 0.00025, r_prod_ = 0.0001, r_deg_ = 0.01, n = 1, d = 10000. There is a ridge of high mean odds ratio with uncorrelated allelic expression to the left and constitutive expression of both alleles to the right. Four example traces show a range of odds ratios. Top traces are gene product counts over time. Bottom traces are the binarized values of each allele over time.

Having established a single metric to summarize allelic correlation for a given simulation, we simulated large parameter sweeps across r_add_, r_off_, k, and n to find parameters leading to different values of allelic correlation. When holding all other parameters constant, there was a ridge of high odds ratio in the parameter space defined by r_add_ and r_off_ (Figure 2B). We found identical ridges of odds ratio across different values of k and n (Figures S2, S3A, S3B). The region of parameter space to the left of the ridge (low r_add_/r_off_) consisted of simulations with lower, purely monoallelic expression. Approaching the ridge, allelic odds ratio increases exponentially to intermediate correlation before reaching a peak. These results closely matched results from a (non-leaky) telegraph transcriptional bursting model of two alleles of a single gene, which also showed a ridge of correlated allelic expression in the parameter space defined by r_on_ (this paper did not model interactions between genes, so only used a basal rate of activation) and r_off_ (Larsson et al., 2021).

Notably, we saw several examples where the odds ratio for allelic correlation became undefined or took on a numeric value that did not fit the standard interpretation of an odds ratio. In the region of parameter space corresponding to low r_add_/r_off_, most simulations were undersampled due to low total expression, causing odds ratios < 1, spuriously indicating anticorrelation between alleles. At very high r_add_/r_off_ ratios and simulations approached a constantly high expression state, the odds ratio approached non-numeric values because of zero values in the contingency table. We therefore lumped together non-numeric odds ratios as indicative of constantly high expression. Overall, we found that low r_add_/r_off_ simulations exhibited low allelic correlation, with increasing allelic correlation up to a ridge of r_add_/r_off_, past which constitutively high biallelic expression dominated, indicating that our model is capable of recapitulating a range of allelic correlation values and can distinguish uncorrelated allelic, highly correlated allelic, and fully biallelic expression.

We wondered whether network size and connectivity had an effect on allelic correlation, since network connectivity could lead to more activating interactions for each gene, increasing the chance for both alleles to simultaneously express when concurrently influenced by multiple regulators. Holding simulation parameters and network connectivity constant (at a connectivity of one), the odds ratio slightly decreased with increasing network size (Figures S4A, S4B). The decrease in odds ratio with network size was expected given that our networks were symmetric: any given burst of activity that could lead to correlated allelic expression will affect a smaller fraction of genes in a larger network given constant parameters. Looking across varying r_add_ and r_off_ for networks of order two and five, the overall distribution of the odds ratio is similar but with an overall decreased magnitude for the five node network compared to the two node network (Figure S4C). In a five node network, increasing connectivity from one to four caused an increase in odds ratio with constant parameters, meaning that more regulating genes led to coordinated allelic expression (Figure S4D). As expected, at the same connectivity, self-looping in the network also increased the odds ratio. We additionally found that the “ridge” of odds ratio is shifted leftward with increasing connectivity and with the addition of self-looping (Figure S4E). Thus, high and low allelic correlations are possible in a range of network architectures, and changing the size and especially the connectivity of the network can have a modest effect on allelic correlation.

### Analysis of ‘multi-bursts’ captures the degree of noise transmission in simulated genetic networks

Input-output correlation corresponded closely with allelic correlation in a limited context. To show whether this correspondence held across many parameter values, we developed several metrics for input-output correlation. We looked at bursts involving multiple genes as the result of input-output correlation (Figures S5A, S5B, S5C). When a given gene bursts in our model, it may or may not cause its downstream target to burst. This propensity corresponds to input-output correlation. In turn, *that* downstream target may or may not cause *its* downstream target to burst, and so on. Thus, the bursting of the initial gene can cause a “multi-burst” of subsequent activity. In simulations with low input-output correlation and low noise transmission, these multi-bursts involved few genes and were short-lived. But in simulations with high input-output correlation and high noise transmission, multi-bursts involved more genes, were longer, and often looped back around to the start of the network. In the limiting case of very high input-output correlation, a single burst can cause a multi-burst that lasts the entire length of the simulation. Thus, we reasoned that the mean number of genes in a multi-burst, the mean multi-burst length, and the number of multi-bursts per unit time were adequate measurements of the input-output ratio of a simulation (Figure S5D).

We calculated the above input-output correlation metrics for simulations with the parameter sets and network architectures described above to find conditions leading to different values of input-output correlation. Similarly to allelic correlation, we found that while holding other parameters constant, all input-output correlation metrics increased with increasing r_add_ and decreased with increasing r_off_ (Figures S5E, S5F, S5G). Like allelic correlation, there was a notable ridge in r_add_ r_off_ space for mean multi-burst length and number of multi-bursts. (For the mean number of genes in a multi-burst, there was no such ridge since the number of genes in a single, simulation-long multi-burst is well defined as equal to the network size.) We found similar ridges across different values of k (affinity) and n (cooperativity) (Figures S6, S7). Additionally, we found that increased network size did not change input-output correlation with constant parameters and increased connectivity increased input-output correlation with constant parameters (Figures S8) Overall, all three of the input-output correlation metrics we evaluated showed that low input-output correlation occurs at low r_add_/r_off_ values, with increasing input-output correlation with increasing r_add_/r_off_.

### Allelic odds ratio follows the relative amount of noise transmission across a systematic range of parameter values and network architectures

So far, we have established metrics for allelic and input-output correlation and found that they appeared in the same region of parameter space, at least in the limited number of cases we checked by eye (Figure 1D). We wanted to confirm this correspondence across a more systematic range of parameter space—in principle, it was possible that in some parameter regimes for which there is an increased correlation of input signal to a given gene’s output may also result in both output alleles to burst at an increased rate but still cause no more co-bursting than expected by chance. In such a situation, we would expect no correspondence between allelic and input-output correlation in our measurements. Conversely, it could be that in parameter regimes that lead to high values of input-output correlation, the input would cause both output alleles to burst nearly simultaneously and thus show more co-bursting than expected by chance. Given that bursts tend to occur on a shorter timescale than the dynamics of signals, we reasoned that the latter scenario might be more likely, because relatively slow fluctuations in the signal could lead to both alleles firing when the signal is high and both alleles not firing when the signal is low, which would look (in aggregate) like an allelic correlation.

To confirm that allelic and input-output correlation corresponded across simulation conditions, we began with visual inspections of their distributions across parameter space (Figure 3A). First, we noticed that both measurements were primarily determined by the r_add_ and r_off_ parameters of our model. Further, both metrics followed a similar ridged distribution in the parameter space of r_add_ and r_off_. Both allelic and input-output correlation followed similar distributions when changing k and network connectivity as well (Figures S9A, S9B). To quantitatively measure the correspondence between the two metrics across this parameter space, we scaled the odds ratio and normalized mean genes in a multi-burst both scaled to between 0 and 1 by subtracting each value by the minimum value and dividing by the range (min-max normalization). We plotted the min-max normalized odds ratio and mean genes in a multi-burst against r_add_ for three constant values of r_off_ and fit exponential curves from which we calculated the K_m_ parameter of fit (Figure 3B). We found the fit curves to be similar by eye, but with K_mm_ values at smaller r_add_ for odds ratio than for mean genes in a multi-burst. The offset of K_m_ holds across values of r_off_, indicating that simulations start to have allelic correlation before having high input-output correlation in our models (Figure 3B, dashed lines). Taken together, we concluded that our model indicates that the relative amount of allelic correlation could provide information for the relative amount of input-output correlation, thus enabling the use of existing allelic correlation data to help answer questions regarding noise transmission.

**Figure 3.**
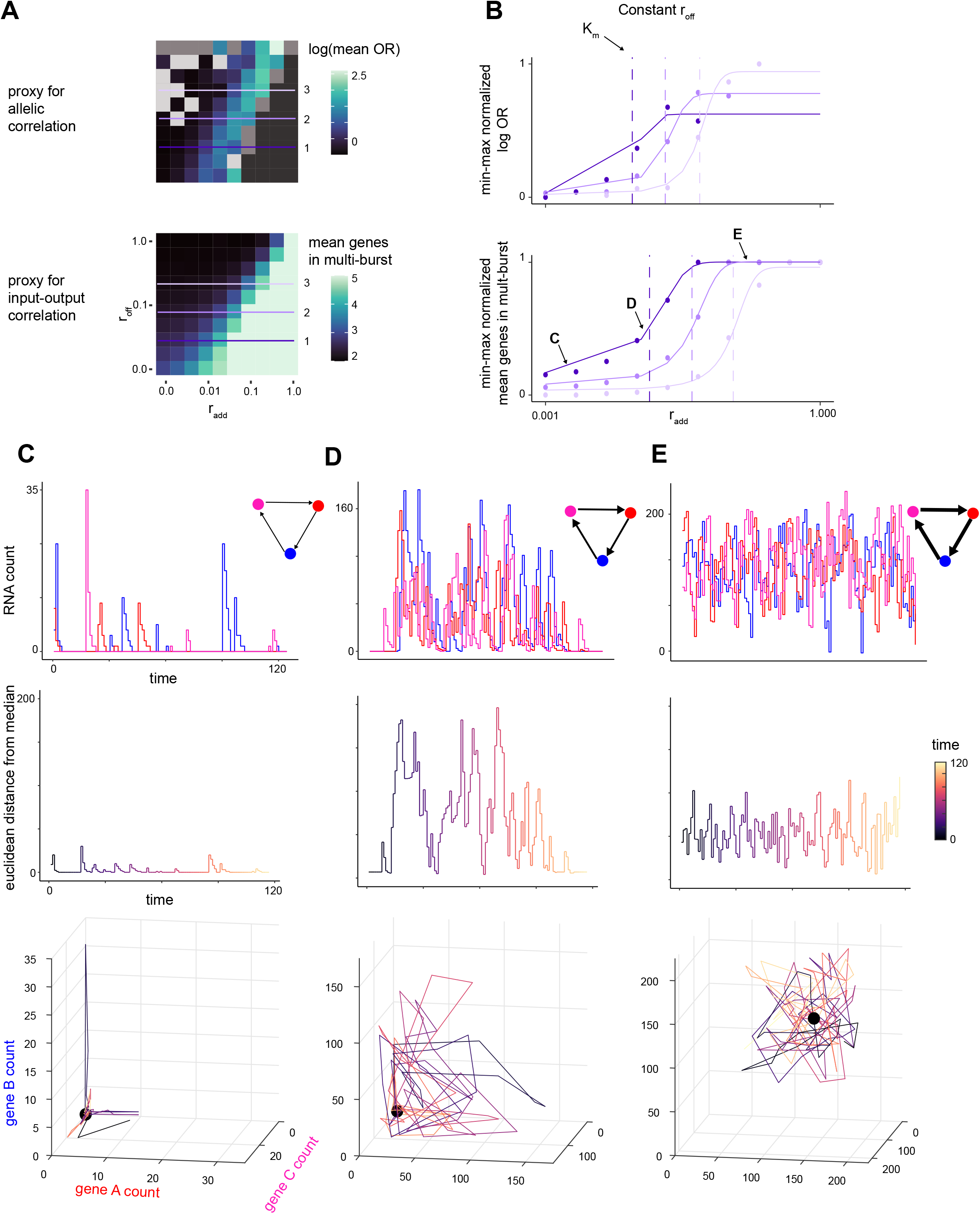
The degree of allelic correlation corresponds with the relative degree of noise transmission across parameter space, marking parameter sets with high variability over time. (A) Comparison of distributions of log transformed mean odds ratio (proxy for allelic correlation) and the mean number of genes in a multi-burst (proxy for input-output correlation) shows similar distributions in r_add_-r_off_ parameter space. (B) The log mean odds ratio and mean genes in burst for three sets of parameters with constant r_off_ (A) fit to similar exponential curves. Log mean odds ratio and mean genes in burst were min-max normalized and fit with (C,D,E) Visualization and analysis of variability for representative simulations from marked regions of the exponential curve. RNA count summed over both alleles for each gene in a three node network, connectivity one network is plotted individually over time (top graph) colored by gene. The euclidean distance from the median is plotted over time (middle graph) with color as the simulation time. The position of the simulation over time is plotted in the three-dimensional gene expression space (bottom graph) is colored by time with median values indicated by the black point. Higher distance over time and longer deviations from the median show that parameter set (D) from the growth part of the curve is more variable over time.

### Parameter regimes with high noise transmission have more frequent and longer correlated deviations from baseline expression

We next were curious what comparative advantages and disadvantages different parameter regimes leading to different levels of input-output correlation may have. Given that noise has been shown to be beneficial in specific contexts (Raj and van Oudenaarden, 2008; Symmons and Raj, 2016), we wondered whether there were potential advantageous functional properties to genetic networks that allowed noise transmission to occur. Specifically, we reasoned that parameter sets which blocked noise transmission (had low input-output correlation) would have lower variability in expression over time but would also be unresponsive to perturbation. Conversely, we predicted that networks with parameter sets that permitted noise transmission (had high input-output correlation) would have more expression variability over time but would be better poised to respond to signals. Does this trade-off between responsiveness and blocking noise transmission exist in our model? To test our above predictions for the variability of simulations with low or high input-output correlation, we picked representative parameter sets from each region of parameter space and looked at variability over time in a three node connectivity of one network (Figures 3C, 3D, 3E). We considered the median mRNA count of each simulation to be the “baseline” expression of each simulation, and so the distance over time from the median captured the deviation from the baseline. Simulations in the part of parameter space that had high input-output and allelic correlation were at further distances from the median for longer times. Another way to capture how far a simulation is from baseline is by plotting the position of the simulation in the expression space defined by the levels of expression of its three genes. We found that simulations in the high input-output correlation region of parameter space are pulled away from the median by an initial bursting of a gene which then transmits to many other genes in the network, causing a long time before the simulation returns to the median. Thus, in simulations with high input-output correlation, a single burst of a gene causes a long-lasting multi-burst that prevents the network from returning to its median for a lengthy period of time.

Conversely, simulations of the part of parameter space with low input-output and allelic correlation had comparatively low expression. When a given gene spontaneously expressed, that expression would rarely transmit downstream to cause a second gene to burst. At the part of parameter space where input-output correlation is the highest (or undefined, depending on the metric used) and allelic odds ratio was undefined, and the expression levels in the simulations were all constantly high, fluctuating about a steady state. Simulations at the extremes of parameter space had comparatively low distance from the median compared to simulations in the middle part of parameter space corresponding to high input-output and allelic correlation. These simulated results suggested that gene regulatory networks in parameter regimes that block noise transmission also would show lower deviation from median expression over time, meaning that such networks would have consistent expression in a basal state. Networks which permit noise transmission would instead frequently deviate far from the median expression over time, leading to longer and often concurrent expression of multiple genes in the network. These long periods of unwanted expression of multiple genes could lead to aberrant downstream responses.

### Noise transmission is required for signal responsiveness but is minimized by trading-off response time

Though high input-output correlation led to potential aberrant downstream responses, these networks were also the ones which most readily transmitted upstream expression variability to activate downstream genes. We reasoned that this observed transmission of random bursts of expression also indicated that high input-output correlation networks were primed for transmission of information. We therefore wondered if high input-output ratio also marked networks that were more responsive to perturbation. To answer this question, we modeled an extrinsic signal input to our network as a node with a constant, high level of RNA (400 molecules) that influenced the activation of a downstream node in the network with a constant r_add-signal_ = 1 (Figure 4A). We initialized the network without the external signal to minimize initialization artifacts. We then ran 100 replicates of introducing the external signal and took the average of the gene expression over time to estimate the final steady state of each gene in presence of the signal. We calculated the dynamic range (dynamic range = log_2_(pre-signal mean expression value / post-signal mean steady state)) for each gene, and the time to reach 0.666 of the post-signal mean for each gene (time constant). We found that the dynamic range in response to signal was maximal in a ridge that overlapped with the ridges of allelic correlation and input-output correlation (Figures 4B, 4C). Both to the left and the right of the ridge, there was little possible dynamic range in response to an external signal. Only the narrow region of parameter space that led to high input-output correlation and thus high noise transmission was poised to respond to external signal.

**Figure 4.**
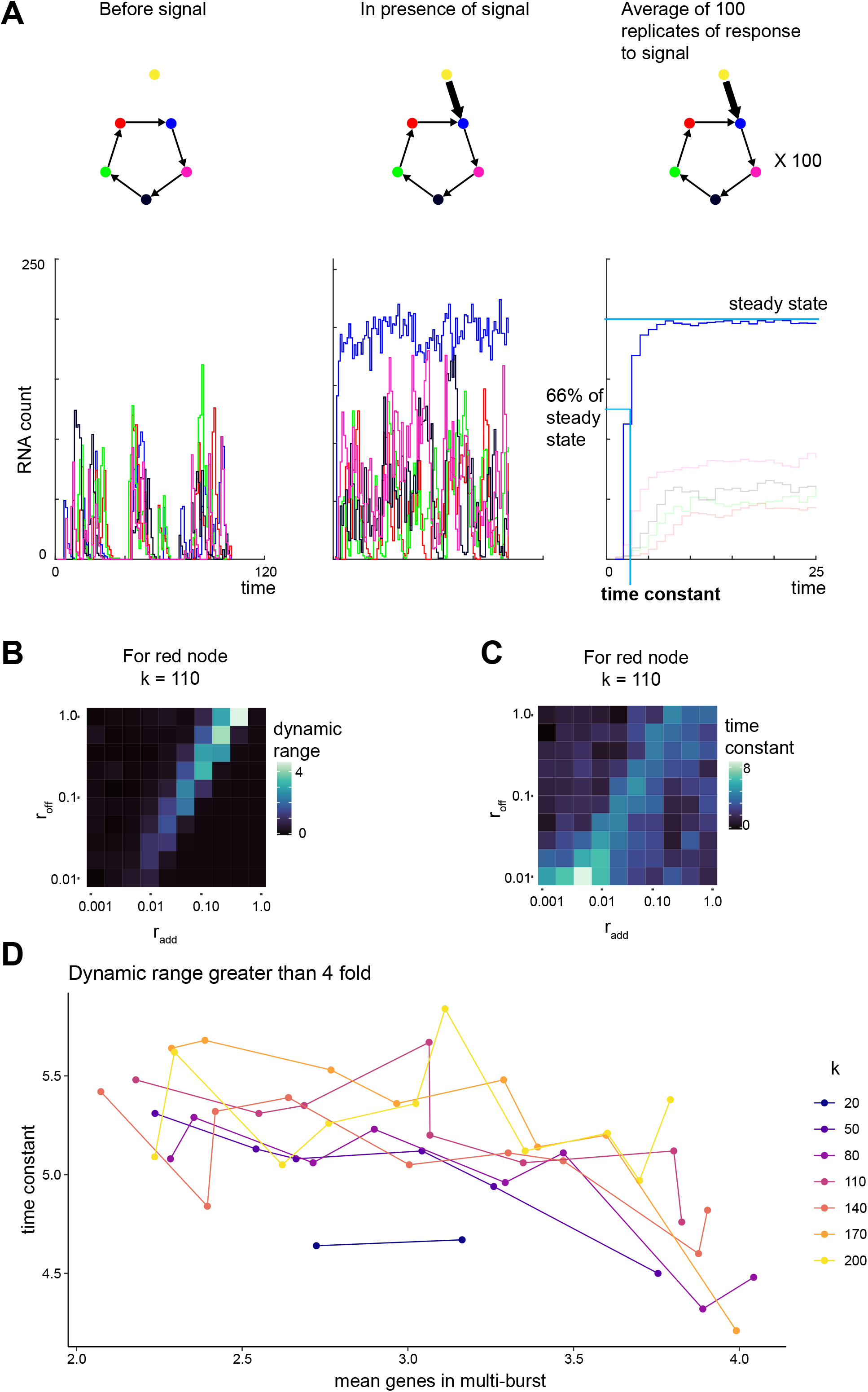
High input-output correlation parameter sets allow response to external signals with a trade-off between response time and baseline noise transmission. (A) Schematic and example traces of signal response simulations. In the absence of an external signal (yellow node), simulations are pre-run to allow initialization of values. Then an external signal is added with strong regulation on a single node of the network. The external signal addition is repeated 100 times. The value of each gene and each time is averaged. From this averaged response, we calculate the steady state and time constant, defined as the time it takes to reach 0.66 of the steady state. (B) Heatmap of dynamic range, defined as the log_2_ fold-change of the post signal steady state to the pre-signal mean value for the red node in the parameter space defined by r_add_ and r_off_ with k = 110 shows a ridge of high dynamic range in the region of high allelic and input-output correlation. (C) Heatmap of the time constant for the same node and same parameter space shows a similar ridge. (D) All parameter sets which show at least a 4 fold response to signal show an inverse relationship between mean total nodes and the time constant across many values of k.

Thus, for a given desired dynamic range to respond to an external signal, there is a minimum noise transmission that is required. We wondered, however, if there could be any other advantage in signal processing from increasing noise transmission beyond the minimum. We hypothesized that further increasing input-output correlation may permit networks to respond more quickly to signal at the cost of higher noise transmission. We tested whether there was a trade-off between the time constant (time to reach 0.666 of post-signal steady state) and basal noise transmission by analyzing simulations which were responsive to signal (log_2_ fold change greater than two). We found that the time constant for such simulations was negatively correlated with the input-output correlation, measured by total genes in a multi-burst (Figure 4D). That is, simulations with higher basal input-output correlation were faster to respond to external signals. This change in response time was modest, with the slowest time constant of 5.84 time units and the fastest with 4.21 time units, but we found that this effect held across values of k. We found that there is a narrow region of parameter space that allows responsiveness to signal in our model, and that within that region, there is a trade-off between response time and basal network fidelity to noise.

### Testing model findings with a published allele-resolved single cell RNA sequencing dataset

#### Genes with high allelic odds ratios are enriched for cell-type specific functions

Armed with the correspondence between allelic correlation and noise transmission, we wondered if we could use sequencing data on allelic correlation to infer the noise transmission properties of genes across the genome. We used a recent single cell RNA sequencing dataset in which the authors performed allele-resolved single cell RNA sequencing on mouse embryonic stem cells and adult fibroblasts (Larsson et al., 2019). Using these data, the authors could infer transcriptional kinetic parameters for each gene. According to their two-state model of transcription, Larsson et al. assumed a parameter space defined by r_on_, equivalent to burst frequency, and r_off_, corresponding to gene inactivation (they did not fit an r_add_ term for additional gene regulation from other genes). We plotted the odds ratio for allelic expression in the parameter space defined by parameters r_on_ and r_off_ estimated from these real data (Figure 5A, supplementary table 1). For ease of visual interpretation, we binned the parameter space (r_on_ was binned into groups of size 0.03 units, and r_off_ was binned into groups of size 0.04 units) and took the mean odds ratio of all genes within that bin, excluding non-numeric odds ratios, which we can then optionally overlay onto the resulting heatmap (Figure 5A, S10). The distributions of allelic odds ratios were similar between embryonic stem cells and adult fibroblasts, and both distributions had a ridge of low allelic odds ratio. We noted that the distributions of allelic odds ratios for embryonic stem cells and fibroblasts appeared to be missing the region of low odds ratio at low r_on_ values compared to the distributions in our models, likely due to technical drop-outs from single cell RNA sequencing.

**Figure 5.**
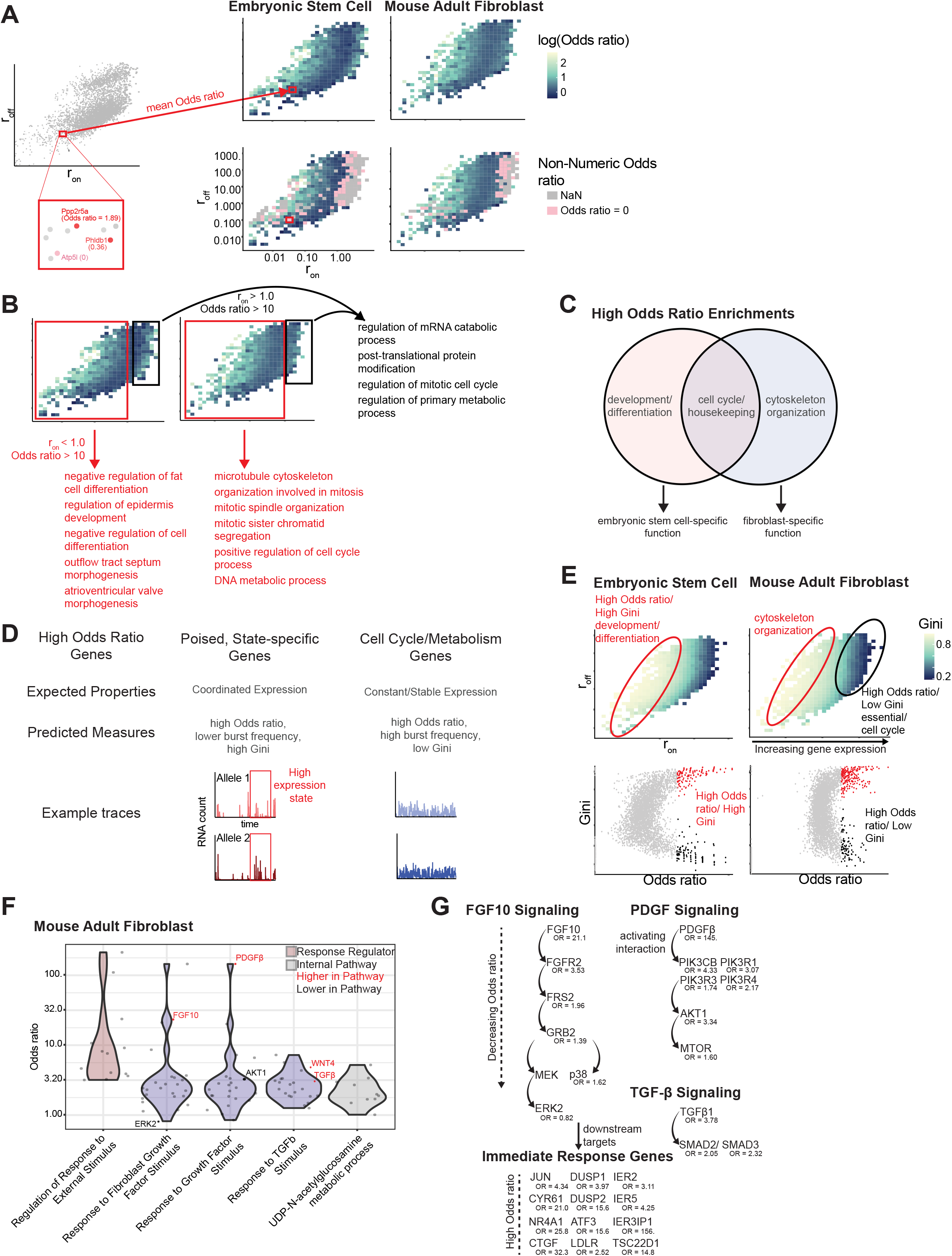
Cell-type specific gene expression identifies functional correlates of model findings. (A) Heatmap displaying allelic odds ratio values in relation to r_on_ (burst frequency) and r_off_ based on allele-resolved single cell RNA sequencing measurements detailed in Larsson, et al. Measurements from mouse embryonic stem cells and mouse adult fibroblasts show a similar distribution of values. (B) Functional annotation, using EnrichR, of distinct groups of genes, identified by their r_on_ and odds ratio values. Regions of high odds ratio (>10) corresponded either to genes with r_on_ > 1, which confer essential “housekeeping” functions, or to r_on_ < 1, which confer cell-type specific functions differing between embryonic stem cells and adult fibroblasts. (C) Schematic diagram displaying the enriched functional categories of genes with allelic odds ratio >10. (D) Schematic diagram displaying the predicted differences between high odds-ratio genes with r_on_ > 1 and r_on_ < 1. Genes with r_on_ < 1 are stipulated to have lower expression and burst frequency on average, but display coordinated high expression states, as indicated by a high Gini coefficient. (E) Correspondence between allelic odds ratio and Gini coefficient. (*Top*) heatmaps displaying Gini coefficient in relation to r_on_ and r_off_. Regions of high odds ratio and low r_on_ occupy a region of parameter space with high Gini coefficient. (*Bottom*) scatterplots relating odds ratio to Gini coefficient, illustrating two distinct groups of genes with high odds ratio: one with high Gini coefficient, and one with low Gini coefficient. (F) Comparison of allelic odds ratio by gene ontology (GO) pathway, specifically comparing genes regulating responses to external signals, individual signaling pathways, and internal pathways. (G) Reconstruction of TGF-b, PGDF, and FGF signaling pathways annotated with allelic odds ratio. Odds ratios strictly and consistently decrease in proximal to distal pathway members.

To find which functional categories were enriched among genes with high odds ratio in each region of parameter space, we performed functional enrichment analysis. We grouped genes according to their odds ratio and r_on_ values, using an odds ratio of ten as a cut-off for low or high odds ratio and likewise a cutoff of one for low or high r_on_. Genes in each category were subsequently subject to functional annotation analysis using EnrichR (Chen et al., 2013; Kuleshov et al., 2016; Xie et al., 2021). The functional enrichment of genes with high allelic odds ratios and high r_on_ values were largely similar between the two cell types and consisted largely of house-keeping genes and genes associated with the cell cycle (e.g. control of transitions between growth and mitotic phases) (Figures 5B, 5C, supplementary tables 2 and 3). One may expect cell cycle genes to have high odds ratios because they are often expressed only at particular parts of the cell cycle, and thus in a population of asynchronous cells there could be a concordance between expression from the two alleles. In contrast, the functional enrichments of genes with high allelic odds ratios and low r_on_ values were not shared between the two cell types. Instead, these gene sets were specific to the function of each particular cell type. In embryonic stem cells, these genes were enriched for functions in differentiation, development, and morphogenesis, but in adult fibroblasts, these genes were enriched for functions including regulation of microtubule cytoskeleton and mitosis. Thus, genes with high odds ratios and low r_on_ values often had functions specific to that cell type. Our model predicts that these cell type-specific genes should be poised to respond to external cues, which may make sense for genes that must perform important cell-type specific functions.

Our analysis appeared to identify two groups of genes with high allelic odds ratio: those with relatively high r_on_ value, which functional annotation analysis appeared to indicate were essential (“housekeeping”) genes, largely common between the two cell types, and those with comparatively lower r_on_ value, which appeared to have cell-type specific function. We expected this latter category of genes to be more prone to burst transmission, and consequently be more likely to show expression only in rare cells (Figure 5D). We sought to capture this rare cell expression using the Gini coefficient as a metric (Jiang et al., 2016; Schuh et al., 2020; Shaffer et al., 2017; Torre et al., 2018). Indeed, genes with low r_on_ and high odds ratio tended to have lower expression and higher Gini coefficient than those with high odds ratio and high r_on_ (Figure 5E, S11, supplementary table 4). Thus, genes with higher odds ratios and low expression tended to be expressed only in rare cells, in agreement with our hypothesis that genes with high allelic odds ratios were more prone to long correlated deviations from baseline.

#### Upstream factors in signaling pathways have higher allelic odds ratios than downstream factors

Our model suggested that genes that need to be responsive to perturbation would have higher noise transmission and thus higher allelic odds ratio. Genes involved in dynamic cellular responses to either external or internal cues may need to be more sensitive to transcriptional regulation by external perturbation. We therefore wondered whether genes involved in externally or internally cued signaling pathways would have high allelic odds ratios. We found that genes encoding proteins that specifically regulate response to external signals had higher allelic odds ratios than all genes encoding proteins involved at all levels of the response to specific signals known to be relevant to fibroblast biology (growth factors such as FGF, TGF, and TGF-b) which in turn had higher allelic odds ratios than UDP-N-acetylglucosamine metabolism, which we considered to be internally regulated and not necessarily an externally cued pathway (Figure 5F, supplementary table 5).

We further found that genes which are upstream in all three pathways had higher allelic odds ratios than their downstream counterparts (Figure 5G). Notably, there was a strict and consistent trend of decreasing odds ratio in more downstream genes. Decreasing allelic odds ratio in more downstream genes suggests that noise transmission was maximal at the start of a signaling pathway but then decreased downstream, which perhaps indicates that the upstream factors in these signaling pathways are tuned to be more responsive to transcriptional regulation.

## DISCUSSION

We demonstrated that a mathematical model of transcription that includes transcription-factor regulation can produce both correlated and uncorrelated allelic expression and that allelic expression correlation closely approximated noise transmission. Parameter sets which led to low noise transmission also produced uncorrelated allelic expression. We also find that some degree of noise transmission is required for a given genetic network to respond to signals, which is also reflected in higher allelic correlation. More noise transmission, however, also leads to greater and longer deviations from basal expression and thus may mimic presence of signal even when no signal exists. To minimize such aberrant network activation while remaining responsive to signals, genetic networks can trade-off response speed to signals. We show that in a recent allele specific single cell RNA-sequencing dataset, genes with high allelic odds ratios are enriched for cell-type specific functions, and that within multiple signaling pathways, genes which are upstream in the pathway have higher allelic odds ratios than downstream genes. Together, our findings suggest that noise transmission, as more easily measured by allelic correlation, must be tightly tuned in genetic networks to allow appropriate signal responsiveness but prevent untriggered activation.

Our work adds to a growing body of evidence suggesting that the autosomal random monoallelic expression observed in allele specific scRNA-seq datasets can be explained by the transcriptional burst hypothesis (Larsson et al., 2021; Rv et al., 2021). However, to date there is no consensus as to any underlying biological advantage that arises from uncorrelated allelic expression. Our findings suggest that uncorrelated allelic expression may very well be a signature of an evolutionary pressure to minimization of noise transmission. We show that networks with high noise transmission (and high allelic correlation) have longer deviations from baseline expression and that these deviations often involve expression of multiple genes. These long, correlated deviations from baseline may in turn cause aberrant activation of downstream cellular processes leading to deleterious effects. Moreover, given the large number of genetic networks and many points of connection between them, if all such networks were to transmit noise, then a random burst would cause a perhaps infinitely long deviation from the basal state. In other words, without some ability to minimize noise transmission, cells would not be able to maintain a basal transcriptional state.

Of course, some genetic networks need to be responsive to perturbation, and our model suggests that such networks must allow a higher degree of noise transmission and thus higher allelic correlation. Genes and genetic networks which require high fidelity but not fast response times should be expected to show low allelic expression correlation. Genes that need to rapidly respond to signals, however, must approach parameter values which lead to higher noise transmission and thus higher allelic expression correlation. Indeed, our analysis of an existing allele specific single-cell RNA-sequencing dataset corroborates these predictions from our model. Genes with high allelic odds ratios and high Gini coefficients were enriched for functions related to cell-type specific signaling pathways, suggesting that genes that encode factors responsible for responding to external signals are poised for response. Our hypothesis was further supported by the striking finding that the allelic odds ratio decreased from upstream to downstream in signaling pathways that respond to external signals. Despite the theoretical downsides to noise transmission as reflected by high allelic odds ratio, our consistent observation of high allelic odds ratios upstream in signaling pathways is consistent with our hypothesis that noise transmission is needed for signal responsiveness.

In our model, once a given network is able to respond to external signals, there is an additional optimization for either signal transmission speed or fidelity to noise in the absence of signal. This trade-off implies that responding quickly comes at the cost of greater aberrant activation due to stochastic, unsignaled bursts. On the other hand, if untriggered activation of a given network were greatly deleterious, then that network may be optimized to respond more slowly in order to more efficiently block transmission of noise. Slow-responding but high fidelity networks may help explain the long timescales of processes that need to be carefully coordinated, such as development (Gregor et al., 2007; Milo et al., 2002). Given the relatively narrow region of parameter space in which the trade-off between basal noise and response speed occurs, emerging technologies to measure multiple RNA species over time in single cells to directly measure noise transmission will be able to more precisely define the limits of these trade-offs (Wan et al., 2021). Alternatively, mathematical models to infer transcriptional noise and dynamics from static measurements may be extended to specifically estimate noise transmission (Gorin and Pachter, 2020; Ham et al., 2021; Komorowski et al., 2011; Munsky et al., 2012; Skinner et al., 2016).

This work suggests that changes in parameters and network architecture lead to changes in signal processing that are intrinsically tied to how genetic networks transmit noise. Network inference studies of gene expression data show that network connectivity can change during state transitions (Moignard et al., 2015; Schlauch et al., 2017). Our results show that at constant parameters, an increase in network connectivity could move a given network from unresponsive to signal to responsive and *vice versa*. Moreover, there is evidence that noise itself may be a tunable parameter of genetic networks. Changes in promoter architecture have been shown to either increase or decrease transcriptional variability and expression timing (Ali and Brewster, 2021; Blake et al., 2006; Jones et al., 2014), providing a tuning method on evolutionary time-scales. On much shorter timescales, epigenetic and DNA repair mechanisms provide another mechanism by which a genetic network can be tuned into or out of signal response regimes (Desai et al., 2021; Weinberger et al., 2012). We posit that transmission of noise is a “necessary evil” of signal responsiveness, and there may be other such tradeoffs as well.

## SUPPLEMENTAL INFORMATION

Supplemental information contains 11 figures and 4 tables

## ACKNOWLEDGEMENTS

We thank the Raj lab members, especially Yogesh Goyal, for scientific discussion and comments on my manuscript. We also thank Amy Azaria for early discussion and pilot analysis in the initial states of the project. A.R. acknowledges NIH Director’s Transformative Research Award R01 GM137425, NIH R01 CA238237, NIH R01 CA232256NIH SPORE P50 CA174523, NIH U01 CA227550, NIH 4DN U01 DK127405. R.H.B. acknowledges support from NIH Training Grant In Computational Genomics T32HG000046 and NIH Medical Scientist Training Program T32GM007170. V.A. acknowledges support from NIH Medical Scientist Training Program 5T32GM007170. L.S. was funded by the Federal Ministry of Education and Research, Germany (Bundesministerium für Bildung und Forschung, BMBF) project TIDY (031L0170B).

## AUTHOR CONTRIBUTIONS

Conceptualization, A.R.; Methodology, R.H.B, L.S., V.A., and A.R..; Software, R.H.B., L.S., V.A., and A.R.; Validation, R.H.B.; Formal Analysis, R.H.B. and V.A.; Resources, A.R.; Investigation, A.R.; Data Curation, R.H.B. and A.R.; Writing - Original Draft, R.H.B.; Writing - Review & Editing, A.R., R.H.B., V.A., and L.S.; Visualization, R.H.B. and V.A.; Supervision, A.R.; Project Administration, A.R.; Funding Acquisition, A.R.

## METHODS

### Network architecture

We use a network model originally developed in Schuh et al. with minimal modifications. Briefly, our model consists of nodes, which represent genes, and directed edges, which represent regulation by another gene. For a network of N nodes, we define an N*N adjacency matrix such that

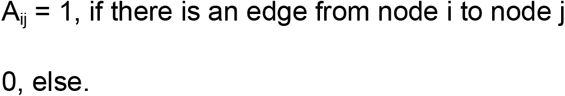

We restrict our networks in three ways. First, we only consider symmetric networks, i.e. networks in which the number of ingoing and outgoing edges within a node and across nodes is identical and either all nodes have a self-loop or not. Second, we constrain our analysis to at least weakly-connected networks. Our final restriction is to only consider non-isomorphic networks, i.e. we discard networks if a bijection exists from the edge space of that network to another in the analysis such that any edge of the first network is projected to the particular edge of the second. Please see Schuh et al. for complete details (Schuh et al., 2020).

These restrictions greatly diminished the number of possible network structures, and allowed us to analyze networks of sizes 2, 3, 4, and 5 nodes which correspond to a total of 24 network architectures.

### Transcriptional bursting model

Our model extends the leaky telegraph model presented in Schuh et al. to include two identical alleles of each gene. In this model, DNA can take on either an active or an inactive state, which translates to a low or a high rate of production of gene products respectively. The inactivation or activation of each allele of a given gene are identical but independent processes. We include interaction terms to the model, where the gene product of an upstream gene influences the rate of DNA activation of the downstream gene dependent on the network structure. We consider a gene product as one mRNA that is faithfully and immediately translated to one protein. We therefore consider the amount of regulation dependent on the mRNA count. The mRNA from each allele are independent but identical: the strength of the regulation depends on the sum of the mRNA from each allele. We model the regulation using the Hill function, given by:

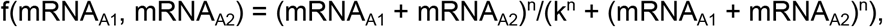

where mRNA_A1_ is the mRNA count of gene A, allele 1, mRNA_A2_ is the mRNA count of gene A, allele 2, n is the Hill coefficient and k is the dissociation constant, n,k >0.

For each allele of each gene, the transitions between inactive and active states along with mRNA production and degradation are modeled by chemical reactions as described previously. Briefly, there are three chemical species: the DNA inactive state, the DNA active state, and the mRNA. The three species interact with each other according to these five chemical reactions:

I -> A

A -> I

I -> I + mRNA

A -> A + mRNA

mRNA -> null.

The chemical reactions are explained by the eight rates/parameters:

| Parameter | Description |
| --- | --- |
| $r_{\text{on}}$ | The rate of DNA activation |
| $r_{\text{off}}$ | The rate of DNA inactivation |
| $r_{\text{prod}}$ | The rate of mRNA synthesis |
| $r_{\text{deg}}$ | The rate of mRNA degradation |
| $r_{\text{add}}$ | Parameter controlling magnitude of additional |
|  | DNA activation rate from gene regulation |
| d | Factor by which mRNA synthesis rate is increased when DNA is in active state |
| k | Dissociation constant of Hill function |
| n | Hill coefficient |

The full model for a single allele which is regulated by gene A, consisting of two alleles A1 and A2, is given below:

| Chemical reaction | Reaction rate | Reaction propensity |
| --- | --- | --- |
| $I \rightarrow A$ | $r_{on} + r_{add} * (mRNA_{A1} + mRNA_{A2})^n / (k^n + (mRNA_{A1} + mRNA_{A2})^n)$ | $(r_{on} + r_{add} * (mRNA_{A1} + mRNA_{A2})^n / (k^n + (mRNA_{A1} + mRNA_{A2})^n)) * I$ |
| $A \rightarrow I$ | $r_{off}$ | $r_{off} * A$ |
| $I \rightarrow I + mRNA$ | $r_{prod}$ | $r_{prod} * I$ |
| $A \rightarrow A + mRNA$ | $d * r_{prod}$ | $d * r_{prod} * A$ |
| $mRNA \rightarrow null$ | $r_{deg}$ | $r_{deg} * mRNA$ |

Where I,A ∈ {0,1}, and I+A=1, and I = 0 marks DNA as in an active state and I = 1 marks DNA as in an active state. mRNA_A1_ and mRNA_A2_ are the mRNA counts of allele 1 of gene A and allele 2 of gene A respectively at the given time, and the parameters are as given above.

In cases where more than one gene influences the expression of a gene, we add the Hill function terms of the respective influencing genes. For example, if the activation of a given allele is influenced by gene A and gene B, both with two alleles 1 and 2, then the rate of the chemical reaction I -> A would be:

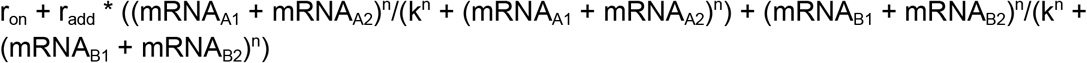

### Parameters

We sought parameter sets which would provide examples of high and low allelic and input-output correlation and then wide ranges of parameters to be able to systematically test correspondence between allelic and input-output correlation. *A priori*, we expected terms in our model that controlled regulation between nodes to be most important for these phenomena. In our model, regulation is at the level of DNA activation, which corresponds to a shift in mRNA steady state from r_prod_/r_deg_ to d * r_prod_/r_deg_. This is controlled by constant term r_on_ and variable term r_add_, which varies according to a Hill function that depends on the count of the regulating mRNA. We wanted activation of DNA that was *not* due to regulation to be rare so that any DNA activation in the presence of a high count of regulating mRNA could reasonably be attributed to regulation. Consequently, we used a constant low value of r_on_ but varied r_add_ as a parameter of interest in our study. Additionally, since the length of each transcriptional burst is exponentially distributed with parameter r_off_, we also varied r_off_ in our study. We also varied k, which serves as the set point for half regulation in the Hill function. In a supplemental parameter set, we varied the Hill coefficient n.

Parameter ensemble one- 700 parameter sets

| Parameter | Values |
| --- | --- |
| $r_{\text{deg}}$ | 0.01 |
| $r_{\text{prod}}$ | 0.0001 |
| $r_{\text{add}}$ | 0.0010, 0.0022, 0.0046, 0.0100, 0.0215, 0.0464, 0.1000, 0.2154, 0.4642, 1.0000 |
| $r_{\text{off}}$ | 0.0100, 0.0167, 0.0278, 0.0464, 0.0774, 0.1292, 0.2154, 0.3594, 0.5995, 1.0000 |
| $r_{\text{on}}$ | 0.00025 |
| d | 100000 |
| k | 20, 50, 80, 110, 140, 170, 200 |
| n | 1 |

Parameter ensemble two (supplemental data only)- 500 parameter sets

| Parameter | Values |
| --- | --- |
| $r_{deg}$ | 0.01 |
| $r_{prod}$ | 0.0001 |
| $r_{add}$ | 0.0010, 0.0022, 0.0046, 0.0100, 0.0215, 0.0464, 0.1000, 0.2154, 0.4642, 1.0000 |
| $r_{off}$ | 0.0100, 0.0167, 0.0278, 0.0464, 0.0774, 0.1292, 0.2154, 0.3594, 0.5995, 1.0000 |
| $r_{on}$ | 0.00025 |
| d | 100000 |
| k | 10 |
| n | 0.2, 0.5, 1.0, 2.0, 4.0 |

### Simulations

We used Gillespie’s next reaction method to simulate our transcriptional bursting model across parameter ensemble one and two (a total of 1200 parameter sets) and 24 network architectures. To adequately sample simulations with low expression, we simulated 20 million time units for Gillespie simulations used to calculate summary statistics. For more manageable file sizes, we used separate 1 million time unit simulations to pull examples of RNA traces. Additionally, to keep file sizes small, we ran parameter ensembles one and two separately. Initialization conditions (t=0) for all simulations was for all DNA to be inactive and all mRNA counts to be 0 molecules. Though the initialization conditions are arbitrary, we allow mRNA counts to reach steady state before visualization and analysis (see below). We performed all simulations in MATLAB 2019a and 2020a.

### Odds ratio analysis

For each simulation, we calculated the odds ratio for simultaneous expression from both alleles of each gene. To allow for each simulation to reach steady state, we first trim the first 100 time units. We then binarize the expression of each individual allele based on whether it exceeds a threshold of 3 molecules to exclude counting small amounts of promoter leak. For each gene, we then use the matlab function *crosstab* to calculate the contingency table for the binarized expression over time of both alleles of the gene. We then calculate the odds ratio for each gene using the MATLAB function fishertest if the resulting contingency table is 2×2; in cases where the contingency table is not 2×2 (such as when two or more values are zero), we assign the odds ratio to be NaN. Since all networks are symmetric with the same parameters for all genes, all genes should have the same odds ratio and so we take the mean of the odds ratio from each gene. We export the data to R for visualization.

### Multi-burst analysis

We analyzed “multi-bursts” for each simulation by first trimming the initial 100 time units. We then sum the gene product counts from both alleles at each timepoint. Next we binarize the expression of each gene based on whether the gene product count exceeds 3 molecules and count the number of genes that are above that threshold at each timepoint. We then find all distinct time intervals in which at one or more time points, two or more genes are expressed above the threshold. These are multi-bursts. For each multi-burst, we calculate the number of time units the multi-burst lasts and the number of genes that exceed the expression threshold during the burst. We then calculate the mean of the burst length and number of genes involved in the burst for across all bursts in the simulation. We also count the total number of bursts in the simulation. We export the data to R for visualization.

### Distance analysis

We calculated distance from the median for each simulation by trimming the first 100 time units and summing the gene product counts from both alleles at each timepoint for each gene. We then calculated the median gene product count for each gene over the length of the simulation. For each gene at each timepoint, we took the square of the difference between the median gene product count and the gene product at that time point. We exported the data to R for visualization

### External signal network and simulations

We extended our network model described above to include one additional node that could be independently modulated to simulate the effect of external signals on the rest of the network. We began with a 5 node, degree 1 network and added a 6th node with a directed edge to node 1. Since we consider this 6th node as an external signal, all chemical reactions for that node were discarded from the model and the gene product count was initialized to either 0 or 400 molecules. The value of 400 molecules was chosen to be much greater than any k value used in the simulations to exert a strong regulation on the network from the signal. In the chemical reaction for gene 1, the signal was given a separate term given by

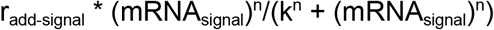

where r_add-signal_ = 1 and mRNA = 0 or 400. Notably, the degradation reaction for the signal node was discarded, so the amount of signal mRNA was constant over time.

Since we are interested in the immediate response dynamics to signal, our strategy for other analyses to arbitrarily set initialization conditions and then only analyze data subsequent to trimming would not be appropriate. Instead, we opted to “pre-run” the simulation without the presence of the external signal (gene product count of node 6 = 0) and then use the conditions at the end of the pre-run to initialize a simulation that included an external signal. Since any given end conditions may not be representative of the steady state of a given simulation, we simulated 100 pre-runs to use as initialization conditions. For each initialization condition from the 100 pre-runs, we then ran 100 replicate simulations with the gene product count of node 6 set to 400 molecules for a total of 10,000 replicates in the presence of signal for each parameter set.

### Dynamic range and time constant calculations

For each parameter set, we want to calculate the change in mean expression that results from stimulation with the signal. In the absence of a signal, we sum the gene product counts of each allele of each gene. We find the mean expression value of each gene before signal addition by taking the mean value of the gene product count for each gene over time, after trimming the first 100 time units (to account for initialization time). Since we perform 100 replicates of the pre-run (no signal), we have 100 mean expression values for each gene for each parameter set. Each of the 100 pre-run replicates maps to 100 post-signal replicates; we took the mean value for each gene (after summing alleles) at each time point over the 100 post-signal replicates that correspond to a given pre-run. The resulting matrix represents the mean value at each time point after exposure to signal (see Figure 4D). Using the matrix, we find the mean value over time of each gene. This is approximately equal to the maximum steady state value in response to the signal since the time to reach the steady state is small in comparison to the length of the simulation (∼5 time units v. 200 time units). We then calculate the time constant, i.e. it takes for each gene to reach 0.666 of its maximum (steady-state) value. If the starting value pre-signal is greater than 0.666 of its max value, we set these time constants to 0. We export the data to R for further analysis and visualization.

In R, we take the mean expression of each gene across all 100 pre-run replicates, resulting in a single pre-signal mean expression for each gene in each parameter set. We do likewise and take the mean of the steady state of each gene across all 100 signal replicates. We then calculate the dynamic range as

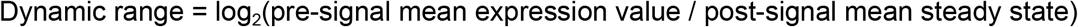

### Functional Enrichment analysis

A dataset (Larsson et al., 2019) comprising allele-resolved single cell RNA sequencing data from 188 embryonic stem cells and 224 mouse adult fibroblasts, along with kinetic parameters for genes comprised therein, was used for functional enrichment analysis. Data were binarized, with each cell being considered to express or not express a given gene, for each allele. Allelic odds ratios were subsequently calculated for each gene.

Gini coefficients were also computed for each gene(Jiang et al., 2016), using Reads Per Kilobase of Transcript (RPKM). In this analysis, a Gini coefficient of 0 implies that for a gene, all cells within a population have the expression. A Gini coefficient of 1 suggests a completely unequal distribution of gene expression that only one cell expresses the gene.. We used the MATLAB function *gini* (Gini coefficient and the Lorentz curve (https://www.mathworks.com/matlabcentral/fileexchange/28080-gini-coefficient-and-the-lorentz-curve/, MATLAB Central File Exchange. Retrieved October 24, 2019.).To relate Gini coefficients, computed on each allele independently, to allelic odds ratio, which summarizes data from both alleles, the average Gini coefficient was used for each gene. Visualizations were performed using R.

Using EnrichR (https://maayanlab.cloud/Enrichr/), functional enrichment analysis was carried out for all genes with allelic odds ratio > 10, odds ratio < 10, odds ratio between 0 and 5, and genes with non-numeric allelic odds ratio. This analysis takes as input lists of genes and identifies among them overrepresented functional categories, as identified by a predetermined list of pathways or “gene ontologies.” Among genes with allelic odds ratio > 10, we subsequently analysed those with r_on_ > 1 and r_on_ <1. In particular, enrichments comprised within the 2021 Gene Ontology Biological Process library were considered.

