## Supplemental figures for "Allelic correlation is a marker of tradeoffs between barriers to transmission of expression variability and signal responsiveness in genetic networks"

#### Supplementary Figure 1

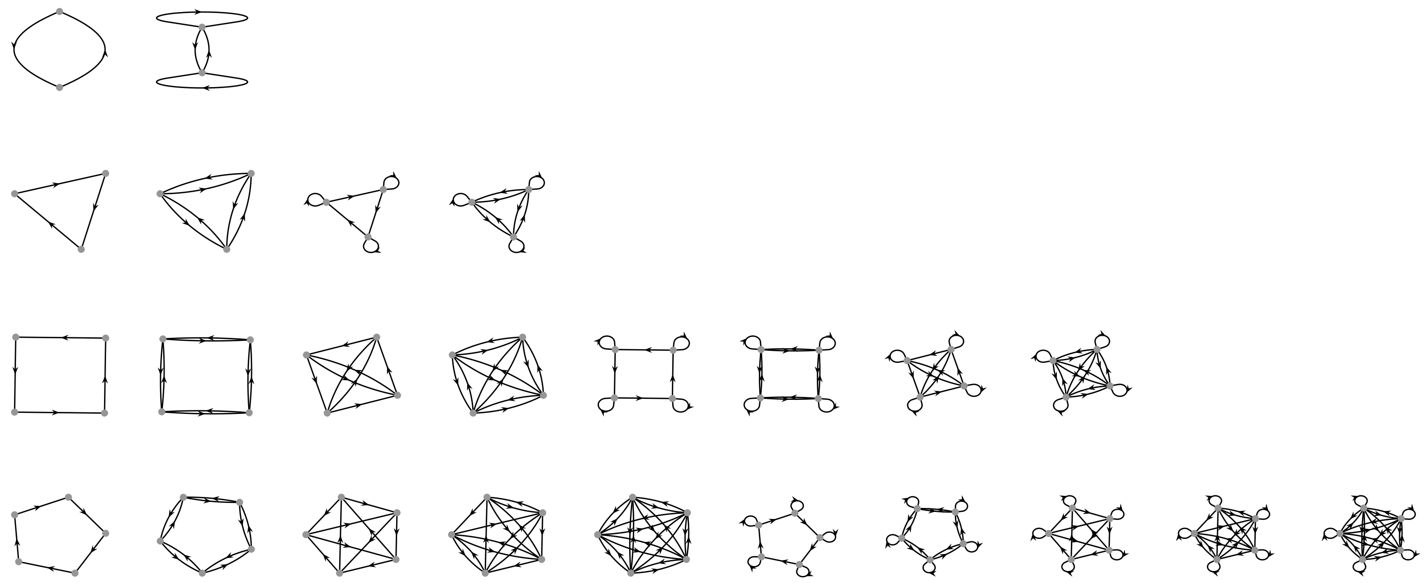

**Supplementary figure 1: Network diagrams of all networks tested.**
Network diagrams with genes as nodes and regulating interactions between genes as directed edges

### Supplementary Figure 2

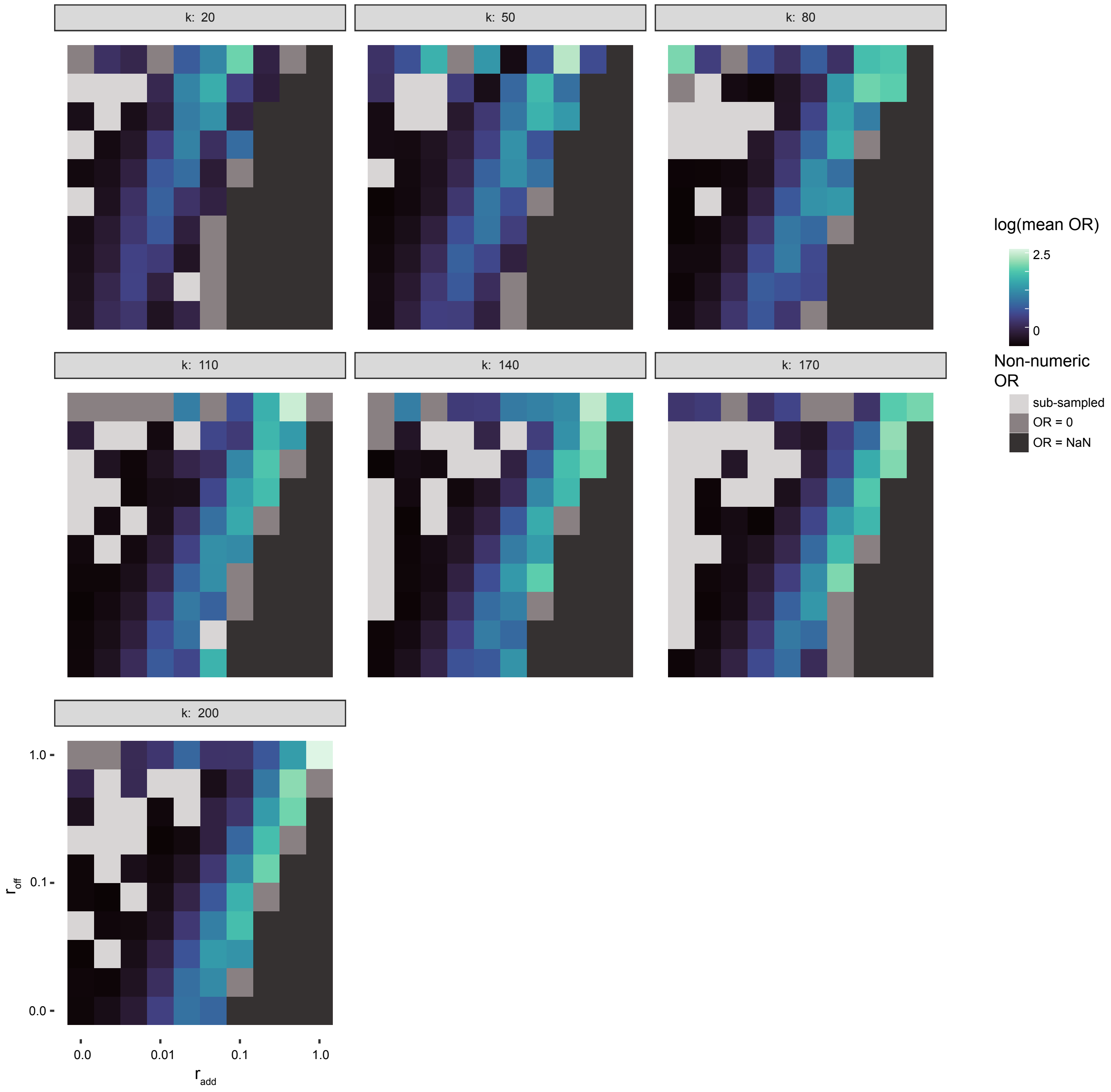


**Supplementary figure 2: Allelic odds ratio shows similar distributions across k values**
Heatmaps of allelic odds ratios across values of k. The distributions are similar to the k = 110 parameter set analyzed in the main figures

### Supplementary Figure 3

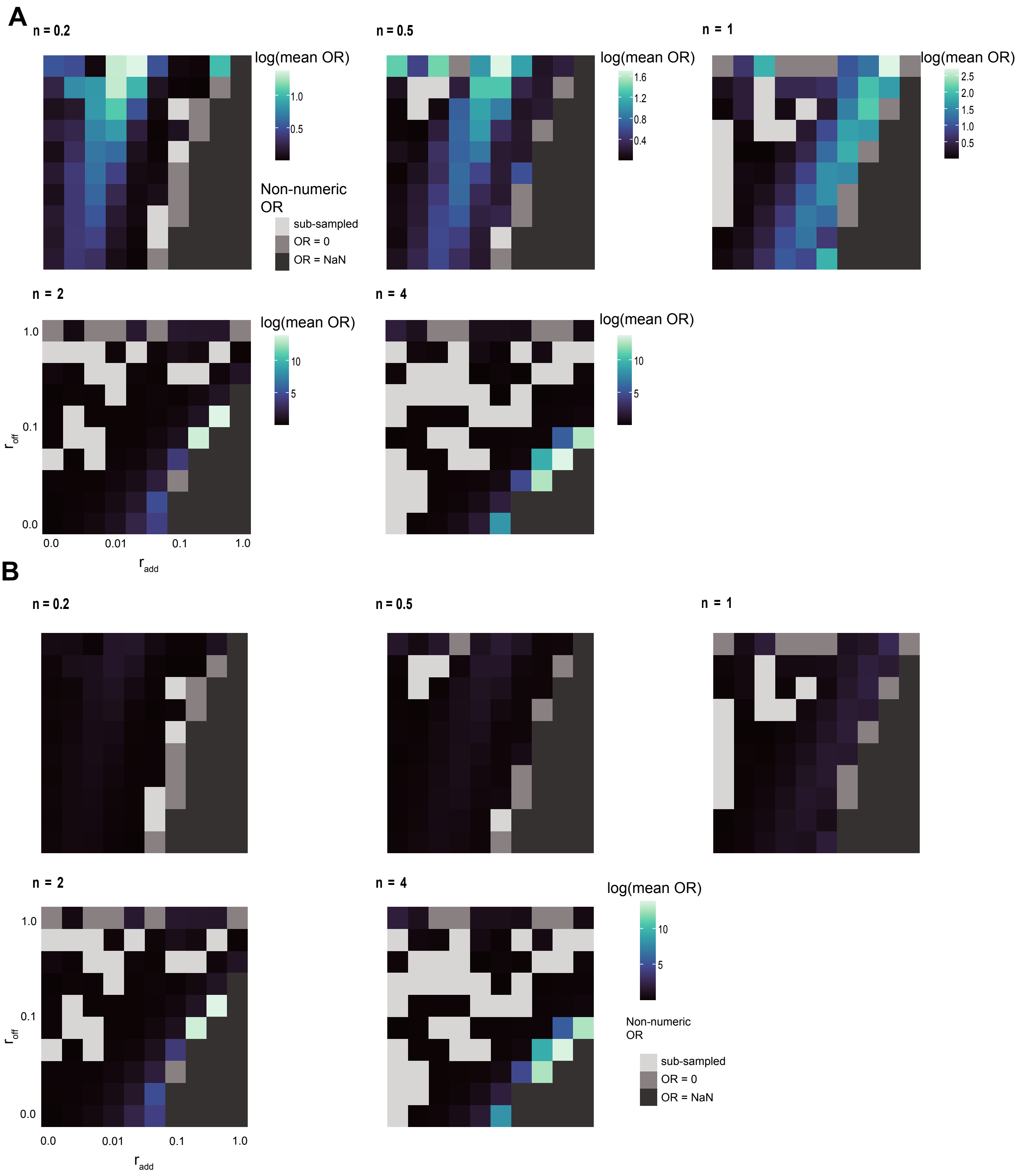


**Supplementary figure 3: Allelic odds ratio shows a ridge of high values across different values of n, but the position dramatically changes**
A. Heatmaps of allelic odds ratio across values of n with individual color scales for each heatmap. The distributions show a similar ridge to the n=1 dataset analyzed in the main figures, but the position is dramatically shifted according to n.
B. The same heatmaps as A but with a single color scale for all heatmaps, emphasizing that the magnitude of allelic odds ratio changes with changes in n.

### Supplementary Figure 4

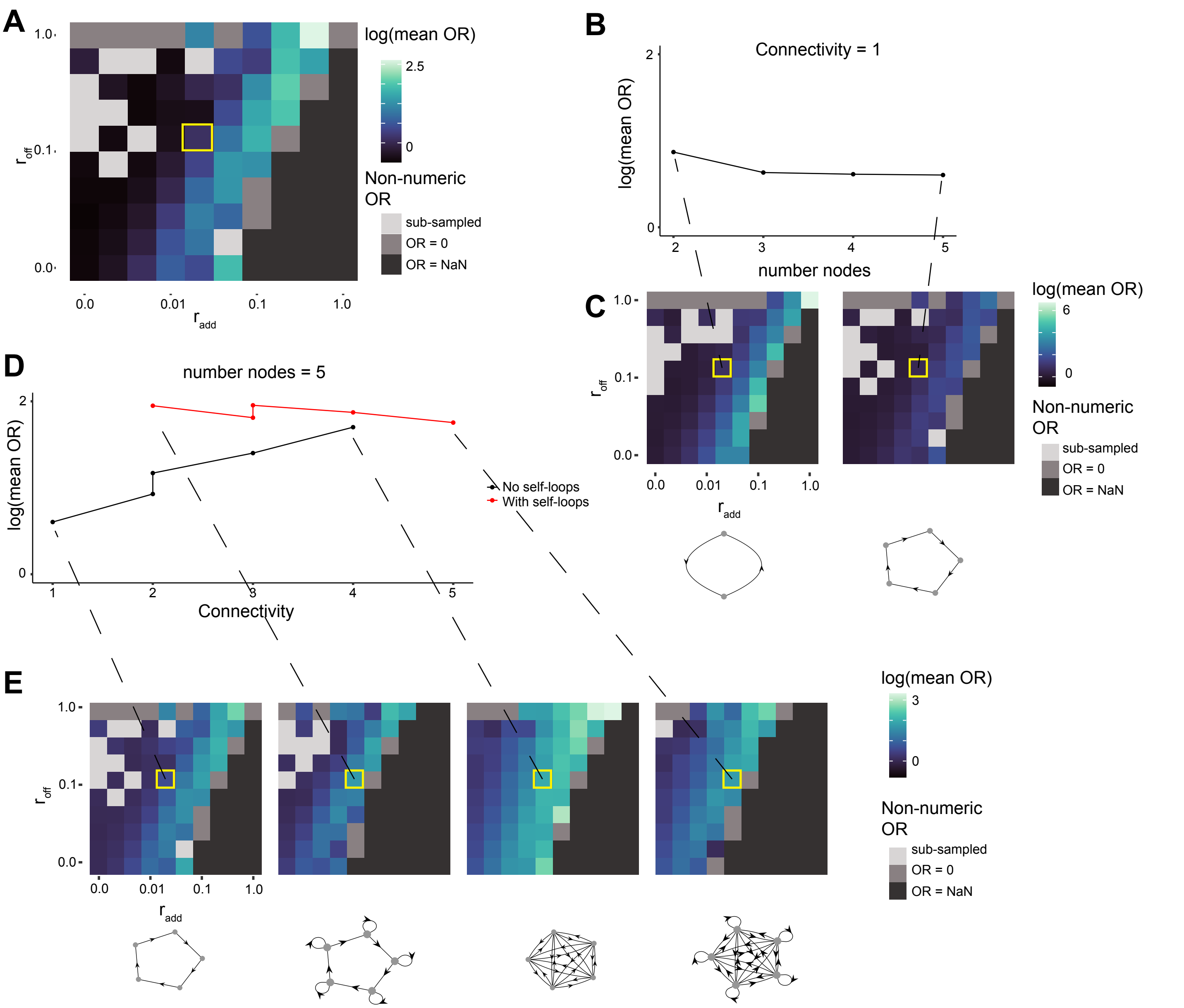


**Supplementary figure 4: Allelic odds ratio decreases slightly with increased network size and increases with connectivity and self-looping**

A. Heatmap of allelic odds ratio in $r_{add}/r_{off}$ parameter space (same as Figure 2). A single parameter set in yellow box is highlighted to hold constant when changing network architecture.

B. Linegraph of log-scaled mean allelic odds ratio against network size (constant connectivity of 1) for the constant parameter set in the yellow box in A. Allelic odds ratio slightly decreases with network size.

C. Heatmaps of entire $r_{add}/r_{off}$ parameter space for network size two and five for context with highlighted parameter set. Network graphs below.

D. Linegraph of log-scaled mean allelic odds ratio against network connectivity (constant number nodes of 5) network for the constant parameter set in the yellow box in A. Networks with self-looping are in red. Allelic odds ratio increases with increased network connectivity. Odds ratio also increases with self-looping.

E. Heatmaps of entire $r_{add}/r_{off}$ parameter space for lowest and highest connectivities with and without self-looping to give context to highlighted parameter set. Network graphs below.

### Supplementary Figure 5

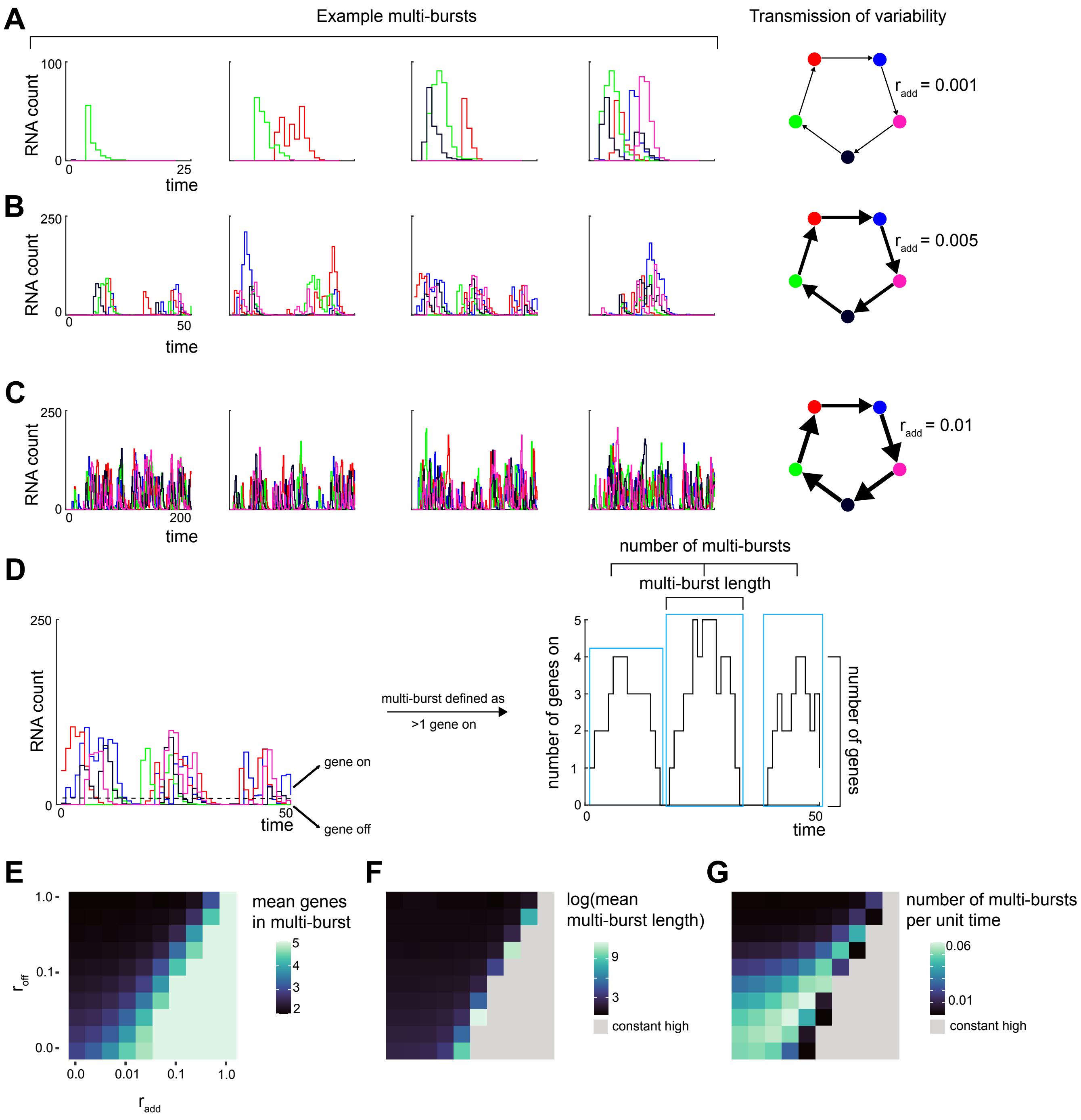


**Supplementary figure 5: Input-output correlation can be measured by quantification of 'multi-bursts'**

A,B,C. Four representative traces of RNA count for three parameter sets with increasing multi-bursts. All other parameters were kept constant besides increasing $r_{add}$ as shown to the right. As $r_{add}$ increases, the length, frequency, and number of genes involved in a multi-burst increases.

D. Schematic showing how multi-bursts are defined and quantified. Gene expression counts (summed over both alleles) are binarized. Runs of gene expression that include at least one time unit of more than one gene on are defined as multi-bursts. We then calculate the number of multi-bursts for each simulation as well as the length of each multi-burst and the number of unique genes involved in each multi-burst.

E. Heatmap of the mean number of unique genes involved in each multi-burst for each simulation. There is a ridge of intermediate values similar to allelic odds ratio.

F. Heatmap of the log-scaled mean length of multi-bursts in each simulation. To the right are simulations which are constantly high, so the mean length is the length of the entire simulation. We color these simulations gray.

G. Heatmap of the number of multi-bursts in each simulation divided by the total length of the simulation.

### Supplementary Figure 6

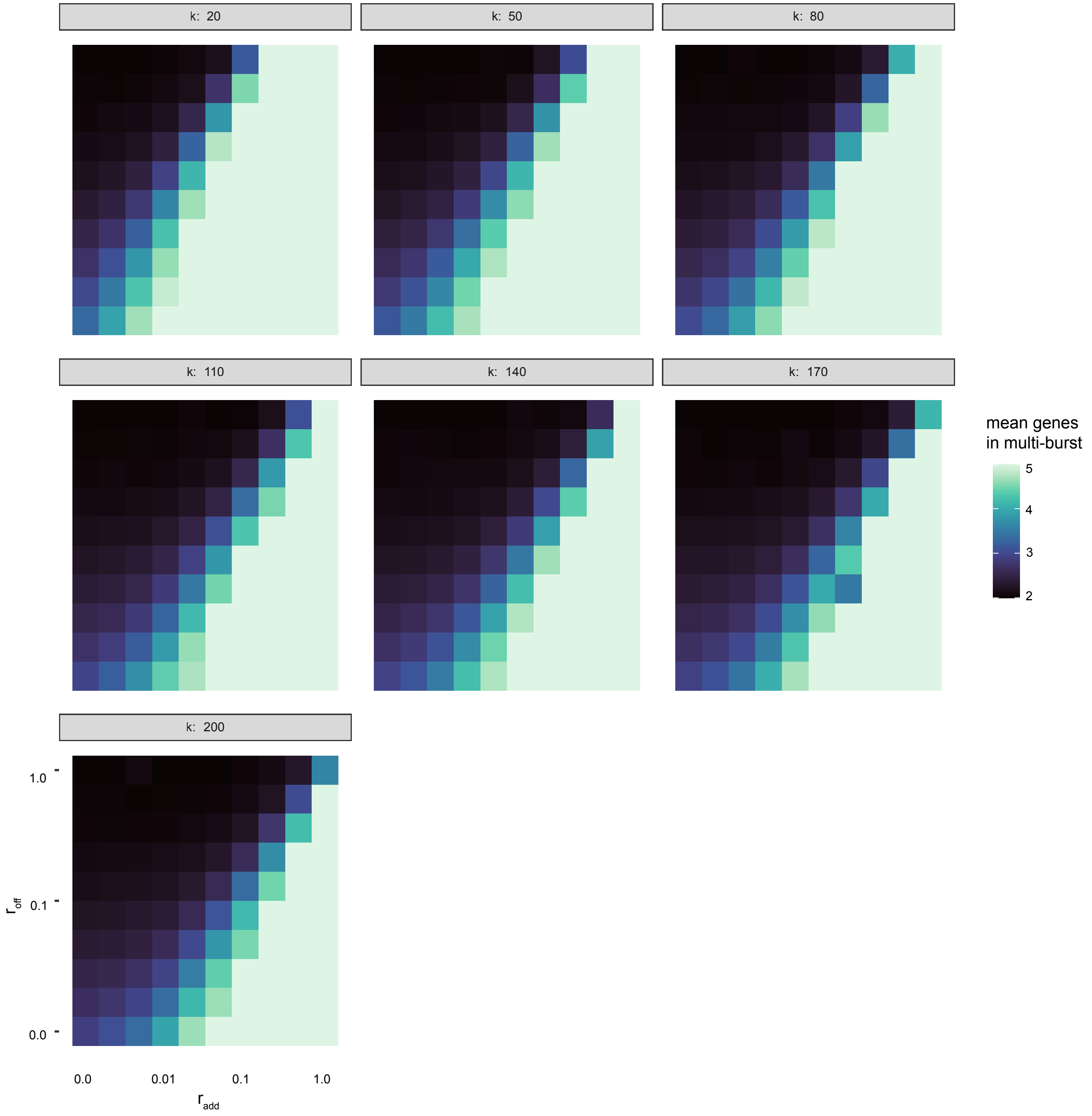


**Supplementary figure 6: Input-output correlation shows similar distributions across k values**
Heatmaps of input-output correlation as measured by mean number of genes in a multi-burst across values of k. The distributions are similar to the k = 110 parameter set analyzed in the main figures

### Supplementary Figure 7

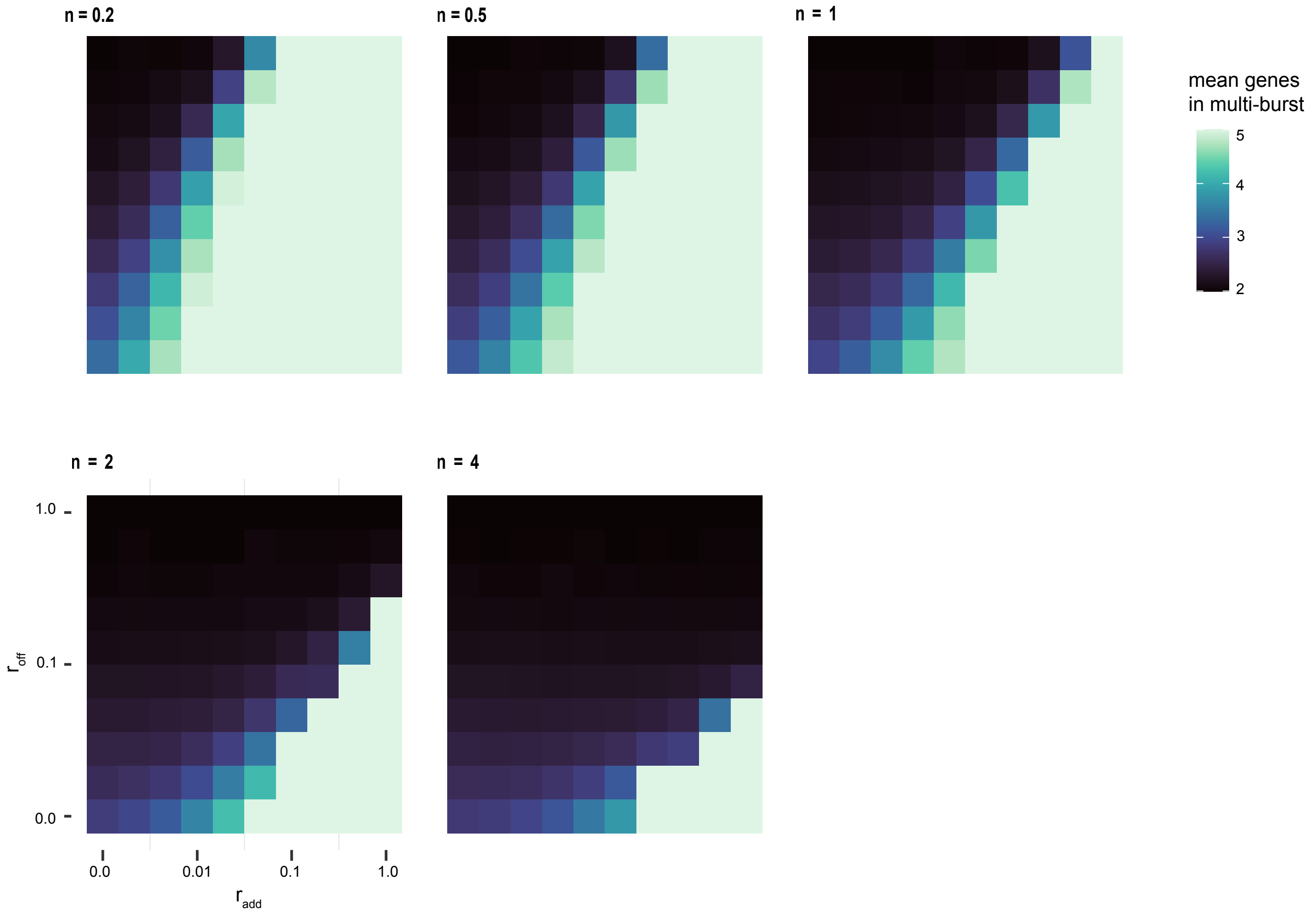


**Supplementary figure 7: Input-output correlation shows similar distributions across n values**
Heatmaps of input-output correlation as measured by mean number of genes in a multi-burst across values of n. The distributions show a similar ridge to the n=1 dataset analyzed in the main figures, but the position is dramatically shifted according to n.

### Supplementary Figure 8

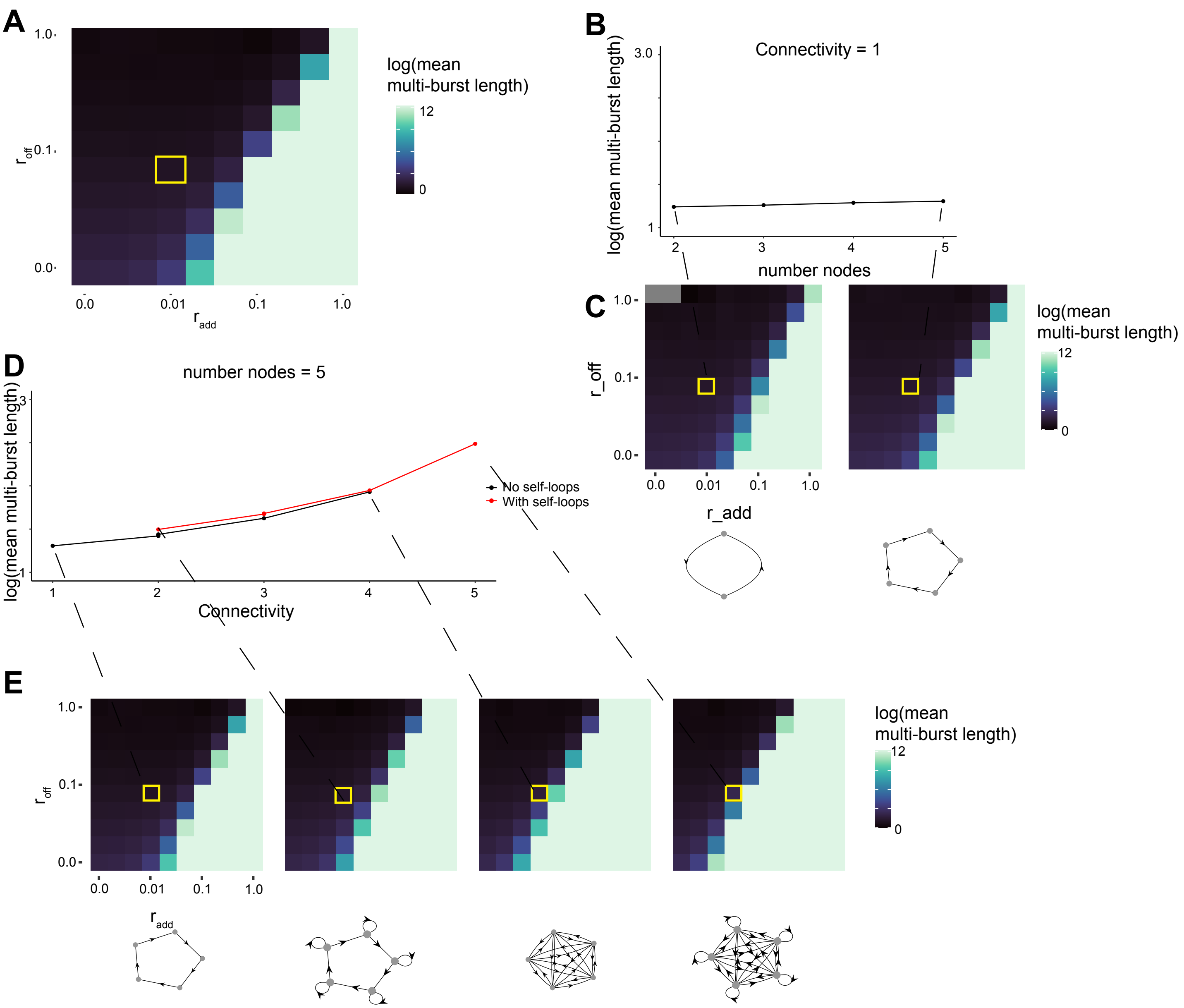


**Supplementary figure 8: Input-output correlation is constant with increased network order and increases with connectivity**

A. Heatmap of input-output odds ratio in $r_{add}/r_{off}$ parameter space (same as Figure 2). A single parameter set in yellow box is highlighted to hold constant when changing network architecture.

B. Linegraph of log-scaled mean multi-burst length as proxy for input-output correlation against network order (constant connectivity of 1) for the constant parameter set in the yellow box in A. Multi-burst length was chosen instead of number of unique genes in multi-burst because the latter metric depends on the total number of nodes in the network. Input-output correlation is constant with increasing network order.

C. Heatmaps of entire $r_{add}/r_{off}$ parameter space for network order two and five for context with highlighted parameter set. Network graphs below.

D. Linegraph of log-scaled mean multi-burst length against network connectivity (constant number nodes of 5) network for the constant parameter set in the yellow box in A. Networks with self-looping are in red. input-output correlation ratio increases with increased network connectivity but is not as dependent on self-looping as allelic odds ratio.

E. Heatmaps of entire $r_{add}/r_{off}$ parameter space for lowest and highest connectivities with and without self-looping to give context to highlighted parameter set. Network graphs below.

### Supplementary Figure 9

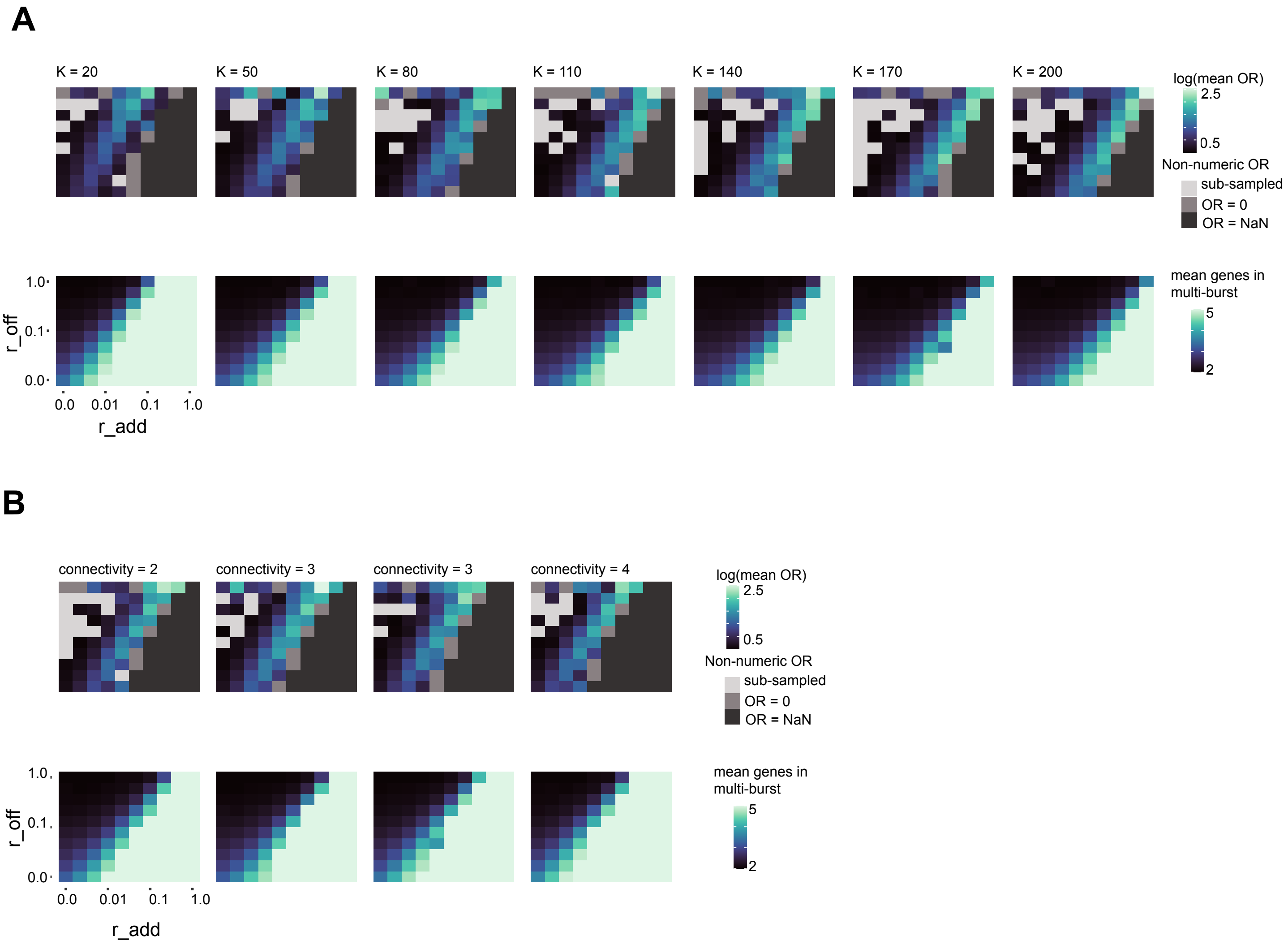


**Supplementary figure 9: Allelic odds ratio and input-output correlation correspond across values of k and network connectivity**
A. Heatmaps of allelic odds ratio (top) and input-output correlation as measured by mean number of genes in a multi-burst (bottom) across values of k. Heatmaps show similar distributions between the two metrics across k values
B. Heatmaps of allelic odds ratio and input-output correlation across network connectivities. Heatmaps show similar distributions across different connectivities.

### Supplementary Figure 10

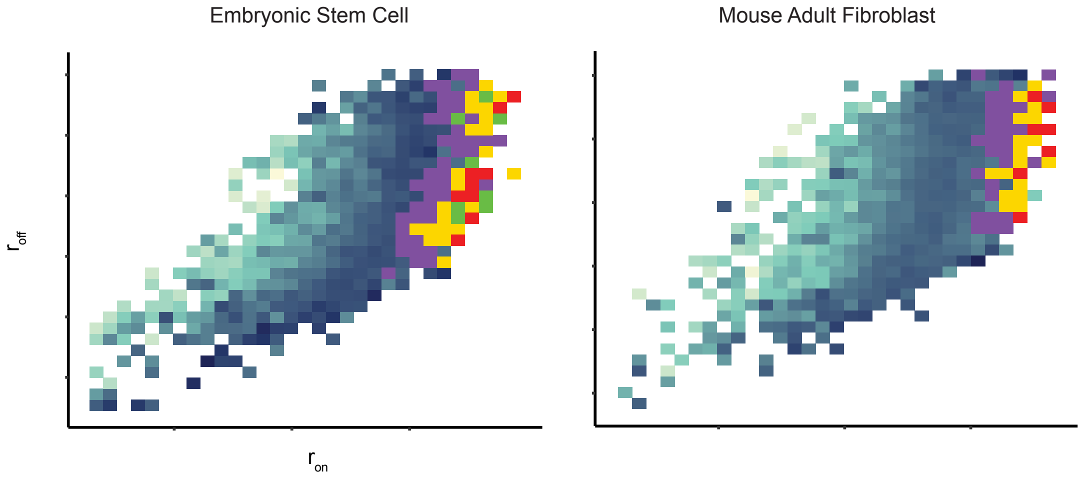


Special Cases of Non-Numeric OR

**Red:** Ubiquitous, biallelic

| Ywhae MAF | Allele C57 Off | Allele C57 On |
|---|---|---|
| Allele CAST Off | 0 | 0 |
| Allele CAST On | 0 | 224 |

OR = 0/0

**Yellow:** Ubiquitous, only one monoallelic possibility

| Fn1 ESC | Allele C57 Off | Allele C57 On |
|---|---|---|
| Allele CAST Off | 0 | 2 |
| Allele CAST On | 0 | 186 |

OR = 0/0

**Purple:** Ubiquitous, monoallelic or biallelic

| Fn1 MAF | Allele C57 Off | Allele C57 On |
|---|---|---|
| Allele CAST Off | 0 | 2 |
| Allele CAST On | 3 | 219 |

OR = 0

**Green:** Not ubiquitous, fully-correlated

| Cox5a ESC | Allele C57 Off | Allele C57 On |
|---|---|---|
| Allele CAST Off | 2 | 0 |
| Allele CAST On | 0 | 186 |

OR = NaN

**Supplementary figure 10: Special cases of allelic odds ratio in allele specific single cell RNA sequencing dataset**
Heatmaps of allelic odds ratio in embryonic stem cells and fibroblasts with the four colors below highlighting bins containing special cases of odds ratios. Red and yellow are ubiquitous expression leading to 0/0 (NaN) values. Purple is ubiquitous expression leading to an OR of zero. Green is fully-correlated leading to a divide by zero (NaN). All of these special cases are enriched at the far right of the heatmaps.

### Supplementary Figure 11

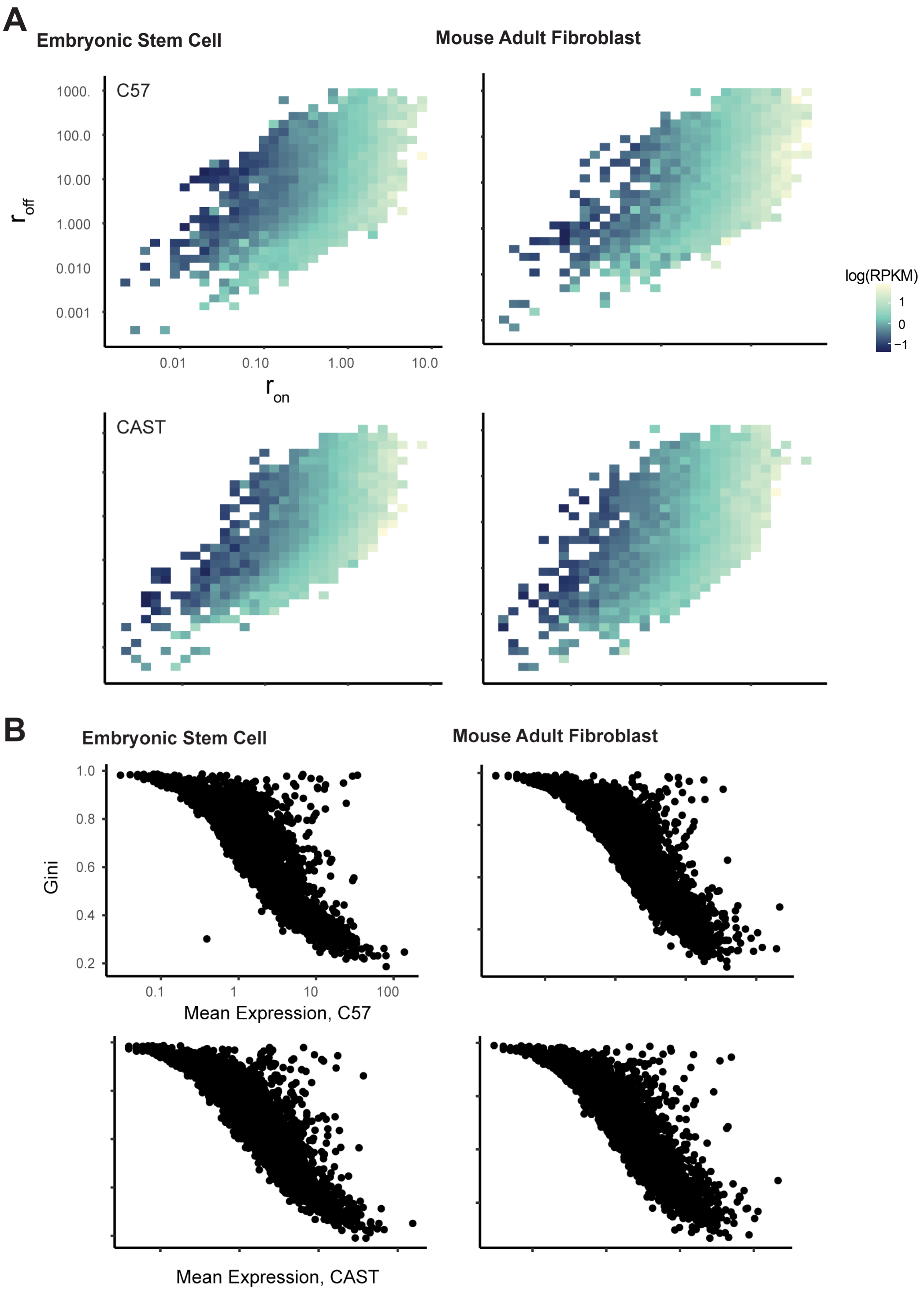


**Supplementary figure 11: Expression in $r_{on}$/$r_{off}$ parameter space shows genes with high allelic ratio ratio and low $r_{on}$ have low expression and high Gini coefficients**

(A) Log transformed RPKM values plotted in $r_{on}$/$r_{off}$ parameter space for embryonic stem cells and adult fibroblasts.

(B) Mean expression plotted against Gini coefficient shows that Gini coefficient is negatively correlated with expression but some genes have high Gini but high expression
